# Recombinant Lloviu virus as a model to study inaccessible zoonotic viruses

**DOI:** 10.1101/2021.08.02.454777

**Authors:** Adam J Hume, Baylee Heiden, Judith Olejnik, Ellen L Suder, Stephen Ross, Whitney A Scoon, Esther Bullitt, Maria Ericsson, Mitchell R White, Jacquelyn Turcinovic, Tran TN Thao, Ryan M Hekman, Joseph E Kaserman, Jessie Huang, Konstantinos-Dionysios Alysandratos, Gabor E Toth, Ferenc Jakab, Darrell N Kotton, Andrew A Wilson, Andrew Emili, Volker Thiel, John H Connor, Gabor Kemenesi, Daniel Cifuentes, Elke Mühlberger

**Affiliations:** Department of Microbiology, Boston University School of Medicine; Boston, USA; National Emerging Infectious Diseases Laboratories, Boston University; Boston, USA; Department of Biochemistry, Boston University School of Medicine; Boston, USA; Department of Physiology & Biophysics, Boston University School of Medicine; Boston, USA; Department of Cell Biology, Harvard Medical School; Boston, USA; Program in Bioinformatics, Boston University; Boston, USA; Institute of Virology and Immunology (IVI); Bern, Switzerland; Department of Infectious Diseases and Pathobiology, Vetsuisse Faculty, University of Bern; Bern, Switzerland; Center for Network Systems Biology, Boston University; Boston, USA; Center for Regenerative Medicine of Boston University and Boston Medical Center; Boston, USA; The Pulmonary Center and Department of Medicine, Boston University School of Medicine; Boston, USA; Department of Pathology & Laboratory Medicine, Boston University School of Medicine, Boston Medical Center; Boston, USA; Institute of Biology, Faculty of Sciences, University of Pécs, Pécs, Hungary; Szentágothai Research Centre, University of Pécs; Pécs, Hungary; Department of Biology, Boston University; Boston, USA

**Author notes:** **Correspondence**, Correspondence and material requests should be addressed to EM. Adam Hume., Elke Muhlberger.

## Abstract

Next generation sequencing has revealed the presence of many RNA viruses in animal reservoir hosts, including many closely related to known human pathogens. Despite their zoonotic potential, many of these viruses remain understudied due to not yet being cultured. While reverse genetic systems can facilitate virus rescue, this is often hindered by missing viral genome ends. A prime example is Lloviu virus (LLOV), an uncultured filovirus that is closely related to the highly pathogenic Ebola virus. Using minigenome systems, we complemented the missing LLOV genomic ends and identified cis-acting elements required for LLOV replication that were lacking in the published sequence. We leveraged these data to generate recombinant full-length LLOV clones and rescue infectious virus. Recombinant LLOV (rLLOV) displays typical filovirus features, as shown by electron microscopy. Known target cells of Ebola virus, including macrophages and hepatocytes, are permissive to rLLOV infection, suggesting that humans could be potential hosts. However, inflammatory responses in human macrophages, a hallmark of Ebola virus disease, are not induced by rLLOV. We also used rLLOV to test antivirals targeting multiple facets of the replication cycle. Rescue of uncultured viruses of pathogenic concern represents a valuable tool in our arsenal against pandemic preparedness.

## Introduction

Zoonotic viruses are a major public health threat. A single spillover event from an animal host into the human population can initiate deadly epidemics or even pandemics. Bats play an important role as natural reservoirs of RNA viruses with the potential to cause significant harm to humans. Examples of bat-borne viruses that have been transmitted to humans, either directly or via intermediate hosts, causing multiple epidemics include Severe Acute Respiratory Syndrome coronavirus (SARS-CoV), Hendra virus, Nipah virus, and Marburg virus (MARV). For other viruses, such as Middle East Respiratory Syndrome coronavirus (MERS-CoV), pandemic SARS-CoV-2, and Ebola virus (EBOV), there is strong evidence that bats might be the natural reservoir, although these viruses have not yet been isolated from bats ^1^. While some of these represent reemerging viruses that were already known to cause severe disease in humans, this list also includes a number of bat-borne viruses that were either unknown or understudied prior to spillover into the human population.

To better prepare for potential future zoonotic epidemics and pandemics, it is necessary to study newly discovered viruses that are closely related to highly pathogenic viruses to both determine their pathogenic potential and to develop and assess potential antiviral therapies. Since 2000, multiple new filoviruses have been discovered via next generation sequencing of samples from wild bats across the globe, including Lloviu virus (LLOV), Bombali virus, and Mengla virus ^2–4^, but the ability to study these viruses has been limited because none of these new viruses have been cultured to date.

One particular virus of concern is LLOV, a filovirus whose viral RNA was initially isolated from carcasses of Schreiber’s bats (*Miniopterus schreibersii*) in Spain and France using deep sequencing and PCR techniques and later found in the same species of bats in Hungary, although no infectious virus has been cultured to date ^2,5,6^. Recently, LLOV re-emerged in bat populations in Northeast Hungary, and again its emergence correlated with unexplained increased mortality among Schreiber’s bats ^5^. Interestingly, many of the bats in which LLOV RNA was found showed symptoms of respiratory infection, but it remains to be determined if LLOV is the causative agent of the disease ^2,5^. One worrying factor regarding the spillover potential for LLOV is the geographic range of the host species, *Miniopterus schreibersii*, which can be found across most of southern Europe, parts of northern Africa, and much of the Middle East, in which over 100 million people live. At the sequence level, LLOV is distinct enough from the ebola- and marburgviruses to be classified within its own genus *Cuevavirus* ^7^. Although the pathogenic potential of LLOV remains unknown, similarities to EBOV and MARV raise concerns that it could be pathogenic for humans. However, the pathogenicity of filoviruses varies considerably, highlighting the need to study LLOV in more detail.

Like many of the genomic sequences of these other uncultured RNA viruses, the LLOV genome is incomplete ^2^. The lack of the terminal genomic sequences that orchestrate viral transcription and replication makes development of a fully wild-type reverse genetic system to rescue the virus impossible. Here we describe the complementation of the LLOV sequence with terminal genomic sequences from other filoviruses in mono- and bicistronic minigenome systems to determine the ability of these sequences to facilitate different aspects of the LLOV replication cycle. Leveraging the data from these complementation assays, we were able to develop a reverse genetic system that facilitated the rescue of infectious recombinant Lloviu virus complemented with sequences from EBOV or MARV (rLLOV_comp_). We analyzed rLLOV_comp_ in various assays, ranging from ultrastructural analysis of infected cells to infection studies with primary human cells, antiviral testing, and host response analysis. Our data suggest that LLOV has the potential to infect humans but does not induce an inflammatory response in human macrophages, a hallmark of severe Ebola virus disease ^8^. The success of this approach provides a roadmap for the rescue and characterization of other uncultured RNA viruses of pathogenic concern for which we currently only have incomplete genomic sequences.

## Results

### Complementation assays to identify functional 5’ genomic ends for LLOV

As a member of the filovirus family, LLOV has a nonsegmented negative sense RNA genome. The published LLOV genome sequence lacks four 3’ terminal nucleotides and almost the entire trailer region. Since the replication and transcription promoter regions of negative sense RNA viruses are contained within the genome ends, called the leader and trailer regions, the lack of the terminal sequences has hampered the generation of recombinant LLOV clones and viral rescue. We previously showed that chimeric LLOV minigenomes that utilize the LLOV leader complemented with four 3’ terminal EBOV nucleotides in combination with the polymerase (L) gene 5’ noncoding region (encoding the 3’ UTR of the L mRNA) and trailer regions (_UTR+tr_) of EBOV, RESTV, and MARV are each replication and transcription competent ^9^. To identify the most potent chimeric trailer region for use in full-length rLLOV_comp_ clones, we generated LLOV minigenomes containing the complete LLOV 5’ noncoding region including the sequence encoding the L 3’ UTR and the 8 known nucleotides of the LLOV trailer complemented with varying lengths of 5’ terminal sequences of the EBOV Mayinga genome (Fig. 1a). Minigenomes with these chimeric LLOV-EBOV 5’ ends were functional, with a general trend of the shorter hybrid trailers being less active (Fig. 1b). Of note, some of these LLOV minigenomes with chimeric trailers were more active than the LLOV minigenome utilizing the full 5’ end of EBOV (EBOV_UTR+tr_, Fig. 1b).

**Fig. 1.**
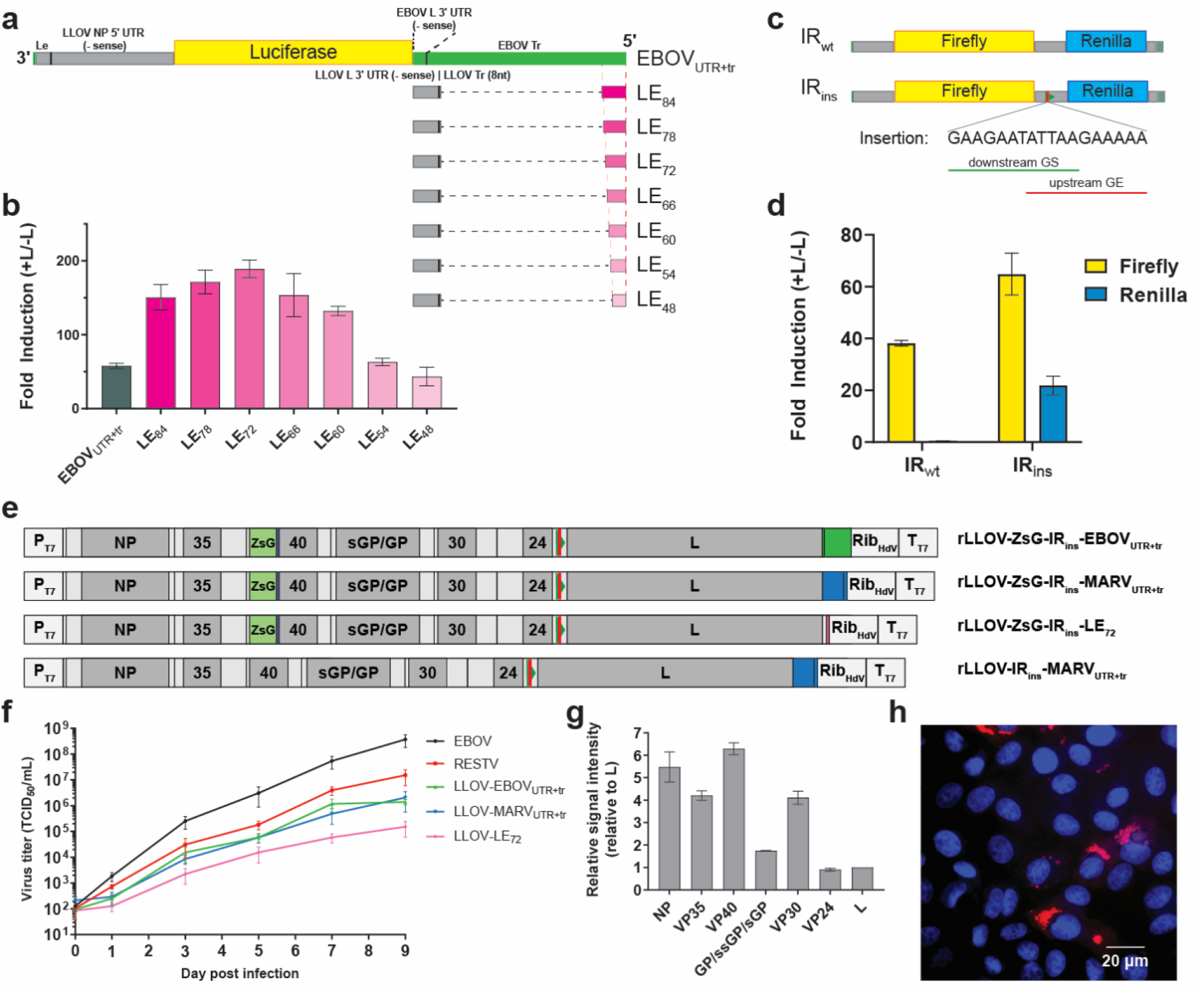
Complementation assays identify sequences that facilitate recombinant LLOV_comp_ rescue. (**a**) Schematic of LLOV minigenomes with EBOV noncoding region (negative sense L gene 3′ UTR) and trailer (top) or hybrid 5′ ends consisting of the LLOV L gene 3’ UTR and trailer complemented with short terminal sequences from the EBOV trailer, designated LEX where X represents the number of added terminal nucleotides from EBOV. Le, leader; NP, nucleoprotein; L, polymerase; Tr, trailer. (**b**) Luciferase-based minigenome assays comparing the 3L5E ^9^ and chimeric 3L5LEX minigenomes. (**c**) Schematic of bicistronic 3L5LE_72_ minigenomes containing firefly and renilla luciferase reporters separated by either the wild-type LLOV VP24-L intergenic region (IR_wt_) or the same intergenic region with an inserted gene border (IR_ins_) consisting of overlapping LLOV gene end (GE, red bar) and gene start (GS, green triangle) signals. (**d**) Luciferase-based minigenome assays comparing the bicistronic minigenomes. (**e**) Schematics of successfully rescued rLLOV_comp_ full-length clones. Noncoding regions are indicated in light gray, LLOV ORFs are in gray, a ZsGreen-P2A reporter (ZsG) is in light green, red bars and green triangles indicate the GE and Gs signals in the IR_ins_ insertion in the VP24-L intergenic region, green indicates EBOV sequences (EBOVUTR_UTR+tr_), pink indicates short EBOV trailer sequences (LE_72_) and blue indicates MARV sequences (MARV_UTR+tr_). rLLOV_comp_ clones are to scale except the T7 RNA polymerase promoter (P_T7_), hepatitis δ ribozyme (Rib_HdV_), T7 RNA polymerase terminator sequences (T_T7_), and IR_ins_, which are enlarged for clarity. (**f**) Growth curve of EBOV, RESTV, and the indicated versions of rLLOV_comp_ in Vero E6 cells. (**g**) Proteomic analysis of LLOV proteins expressed in SuBK12-08 cells infected with rLLOV-ZsG-IR_ins_-EBOV_UTR+tr_ at a multiplicity of infection (MOI) of 1 at two days post-infection (dpi). Signal intensities of viral proteins are plotted relative to L signal intensity within the same sample. (**h**) RNA FISH analysis of Vero E6 cells infected with rLLOV-IR_ins_-MARV_UTR+tr_ at an MOI of 1 at 1 dpi. Red, negative sense (genomic) LLOV RNA clustered in viral inclusions. Cell nuclei were stained with DAPI (blue).

### Full-length LLOV clones containing bicistronic VP24-L genes cannot be rescued

Based on the minigenome data, we successfully generated full-length LLOV clones that contained the EBOV_UTR+tr_, MARV_UTR+tr_, or LE_72_ as 5’ genome ends. LE_72_ genome ends were chosen because this minigenome performed best in the minigenome assay (Fig. 1b). Since there are no commercially available antibodies against LLOV, we also constructed these rLLOV_comp_ full-length clones with the addition of a ZsGreen reporter attached to the VP40 gene via a P2A sequence as previously described for EBOV ^10^. Viral rescue was attempted by transfecting cocultures of African green monkey kidney (Vero E6) and human hepatocarcinoma (Huh7) cells with full-length clone rLLOV_comp_ plasmids along with codon-optimized LLOV support plasmids as previously described for the LLOV minigenome system (Fig. 1e, Supplementary Fig. 1) ^9^. However, repeated attempts to rescue these clones were unsuccessful (Supplementary Fig. 1). One striking observation regarding the published LLOV genomic sequence is the lack of gene end (GE) and gene start (GS) signals within the VP24-L intergenic region (IR), which has led to speculation that these genes may be expressed bicistronically ^2^. However, this would be unique, as there are, to our knowledge, no instances of bicistronic or polycistronic genes for any nonsegmented negative sense RNA viruses outside of the bornavirus family ^11,12^. To test whether this lack of GE and GS signals might play a role in the expression of both the LLOV VP24 and L genes, we constructed and tested a bicistronic minigenome that included the wild-type VP24-L IR between firefly and renilla luciferase reporter genes (Fig. 1c). Although we saw expression of the first reporter gene in a minigenome assay, expression of the second gene was not observed (Fig. 1d). The addition of conserved LLOV GE and GS signals within this IR in the bicistronic minigenome (IR_ins_), however, facilitated efficient expression of both reporter genes, indicating that there is a strict requirement for flanking GS and GE signals for each LLOV gene (Fig. 1d).

### Rescue and characterization of recombinant Lloviu viruses

Incorporation of the VP24-L IR_ins_ sequence into rLLOV_comp_ full-length genomes facilitated the rescue of three different versions of rLLOV_comp_ containing the ZsGreen-P2A-VP40 reporter, EBOV_UTR+tr_, MARV_UTR+tr_, and LE_72_ (Fig. 1e and Supplementary Fig. 1) The ability of these viruses to be rescued only with the addition of the IR_ins_ sequence further highlights the strict functional need for GS and GE sequences within the LLOV genome. This raises questions about whether the published LLOV VP24-L IR sequence is potentially incomplete, possibly due to masking of IR sequences on the viral genome by abundant viral mRNA or due to possible sequencing difficulty caused by RNA secondary structures. However, since the sequencing files from the original discovery of LLOV are not publicly available, these hypotheses cannot be assessed ^2^.

We next sought to compare replication kinetics of rescued Lloviu viruses with recombinant EBOV (Mayinga) and RESTV (Pennsylvania) containing ZsGreen-P2A-VP40 reporters. All three rLloviu viruses grew to lower titers in Vero E6 cells, a cell line frequently used for filovirus replication kinetics, and replicated more slowly than both the rEBOV and rRESTV, with rLLOV-LE_72_ representing the slowest-growing virus (Fig. 1f). Each of the LLOV proteins was detected at abundant levels as determined by proteomic analysis of a bat (*M. schreibersii*) kidney cell line (SuBK12-08) ^13^ infected with rLLOV_comp_, with a trend of viral proteins encoded closer to the 3’ genomic terminus producing higher relative signal intensities, indicative of greater abundance (Fig. 1g, Supplementary Fig. 2). This mirrors the transcriptional gradients observed in EBOV and MARV infections ^14^. The published LLOV sequence contains a hypothetical open reading frame predicted to be encoded on the antigenome overlapping the VP24 gene (rcORF), but proteomic analysis did not indicate any evidence for the expression of this rcORF.

Vero E6 cells infected with rLLOV_comp_ showed typical filoviral inclusions by RNA fluorescence in situ hybridization (FISH) analysis, as visualized by clustered localization of viral genomic RNA (Fig. 1h). Similarly, infection of both human (Huh7) and bat (SuBK12-08) cells with rLLOV_comp_ showed characteristic filovirus inclusions by ultrastructural analysis, with both transverse and longitudinal sections of filamentous nucleocapsids visualized by electron microscopy (EM), similar to filovirus inclusions ^15^ (Fig. 2a-d and Supplementary Fig. 3). Lloviu virions were observed budding from infected cells (Fig. 2c; see Supplementary Fig. 3 for EBOV budding). We also examined rLLOV_comp_ particles by both transmission electron microscopy and cryoelectron microscopy, observing typical filoviral virion morphologies including particles with simple filamentous, 6-shaped, filamentous with terminal membrane protrusions, and branched filamentous structures (Fig. 2f and g), similar to EBOV virion morphologies (Supplementary Fig.3).

**Figure 2.**
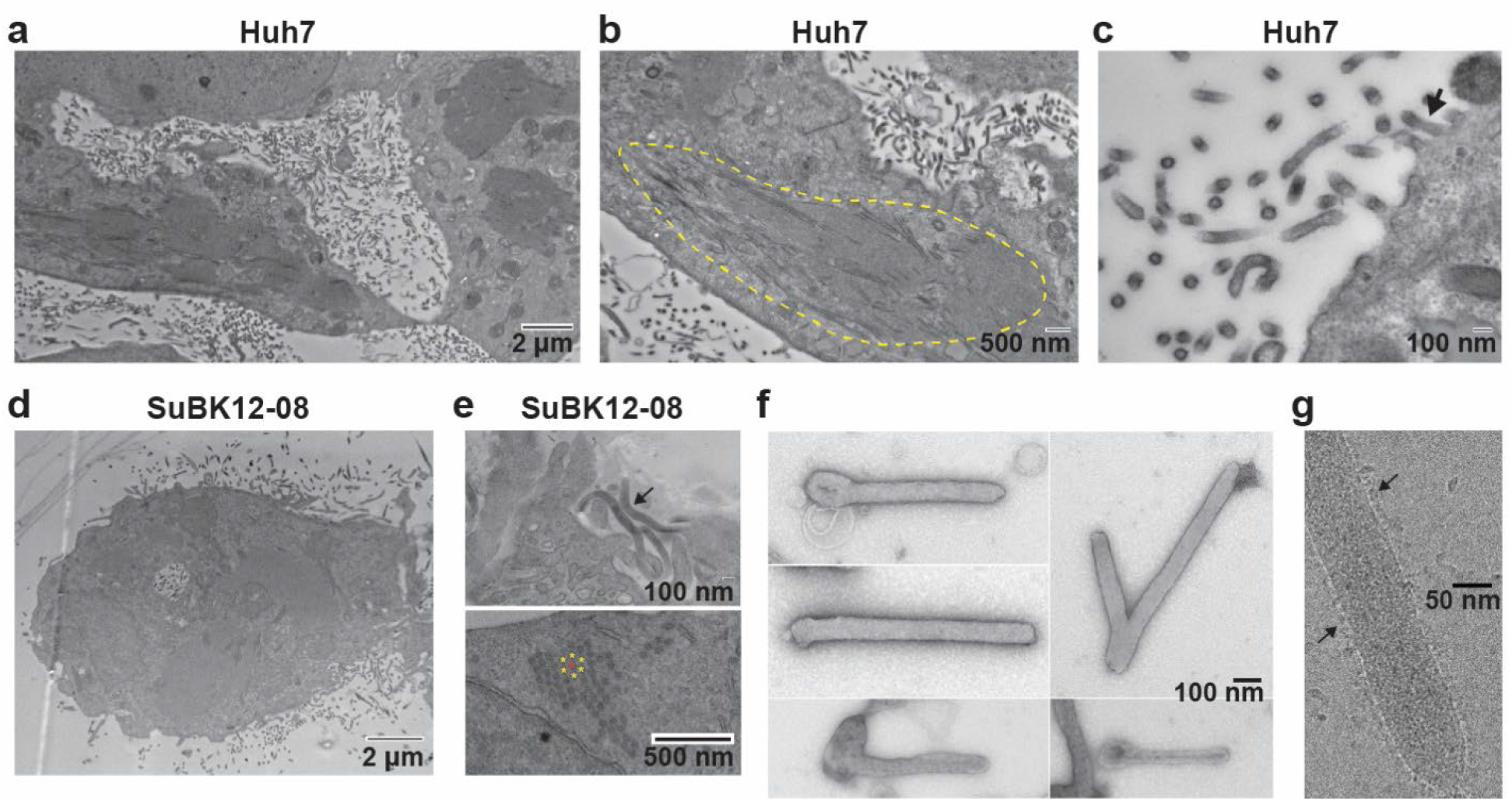
Electron microscopy (EM) of rLLOV_comp_-infected cells and virions. (**a-e**) Transmission EM of Huh7 or SuBK12-08 cells infected with rLLOV-IR_ins_-MARV_UTR+tr_ at an MOI of 1 and fixed at 3 dpi. (**a**) Release of viral particles from infected Huh7 cells. (**b**) Circled area indicates the accumulation of filamentous LLOV nucleocapsids into inclusions. (**c**) Mature virions budding from the cell surface (arrow). (**d)** rLLOV_comp_-infected SuBK12-08 cell releasing viral particles. (**e**) Filamentous viral particles (top panel, arrow) and cross section of viral inclusions (bottom panel). The asterisks indicate the typical honeycomb pattern of filovirus nucleocapsids within the rLLOV_comp_ inclusions. (**f**) EM of negatively stained, isolated rLLOV-IR_ins_-MARV_UTR+tr_ virions. (**g**) CryoEM of rLLOV-IR_ins_-MARV_UTR+tr_ virions. LLOV glycoproteins studding the particle surface are indicated with arrows.

RNA-Seq analysis of the rLLOV_comp_ stocks indicated few changes, with limited single nucleotide variants identified (Supplementary Fig.4). No consensus changes were observed in any of the filovirus proteins that have previously been identified to play a role in antiviral responses or cross-species adaptation, including VP35, VP40, glycoprotein (GP), and VP24 ^16^ (Supplementary Fig. 4). Additionally, after high-throughput cloning and sequencing of the 5’ terminal genomic sequences of the rLLOV-EBOV_UTR+tr_ and -MARV_UTR+tr_ stocks we observed very few changes in the 5’ terminal sequences of either of these viruses. These changes were all detected at low abundance despite good sequencing coverage, with the exception of an untemplated AC dinucleotide at the 5’ terminal end of the rLLOV-EBOV_UTR+tr_ sequence found in about 20% of sequence reads (Supplementary Fig. 5). The relative lack of mutations at the 5’ end of these viruses indicates that the LLOV polymerase has some flexibility in promoter recognition.

To gain insight into LLOV host cell tropism, we compared the ability of rLLOV_comp_ and rEBOV to infect two key target cell types during filovirus infections in humans, macrophages and hepatocytes ^17^. Infection of both monocyte-derived macrophages (MDMs) and induced pluripotent stem cell (iPSC)-derived, primary-like hepatocytes (iHeps) ^18^ with rLLOV and rEBOV revealed similar infection rates, with higher infection rates with rEBOV in the iHeps at later time points (Fig. 3a and b). These data indicate that there is no species or cell-type tropism block that would prevent LLOV from being able to productively infect human cells.

**Figure 3.**
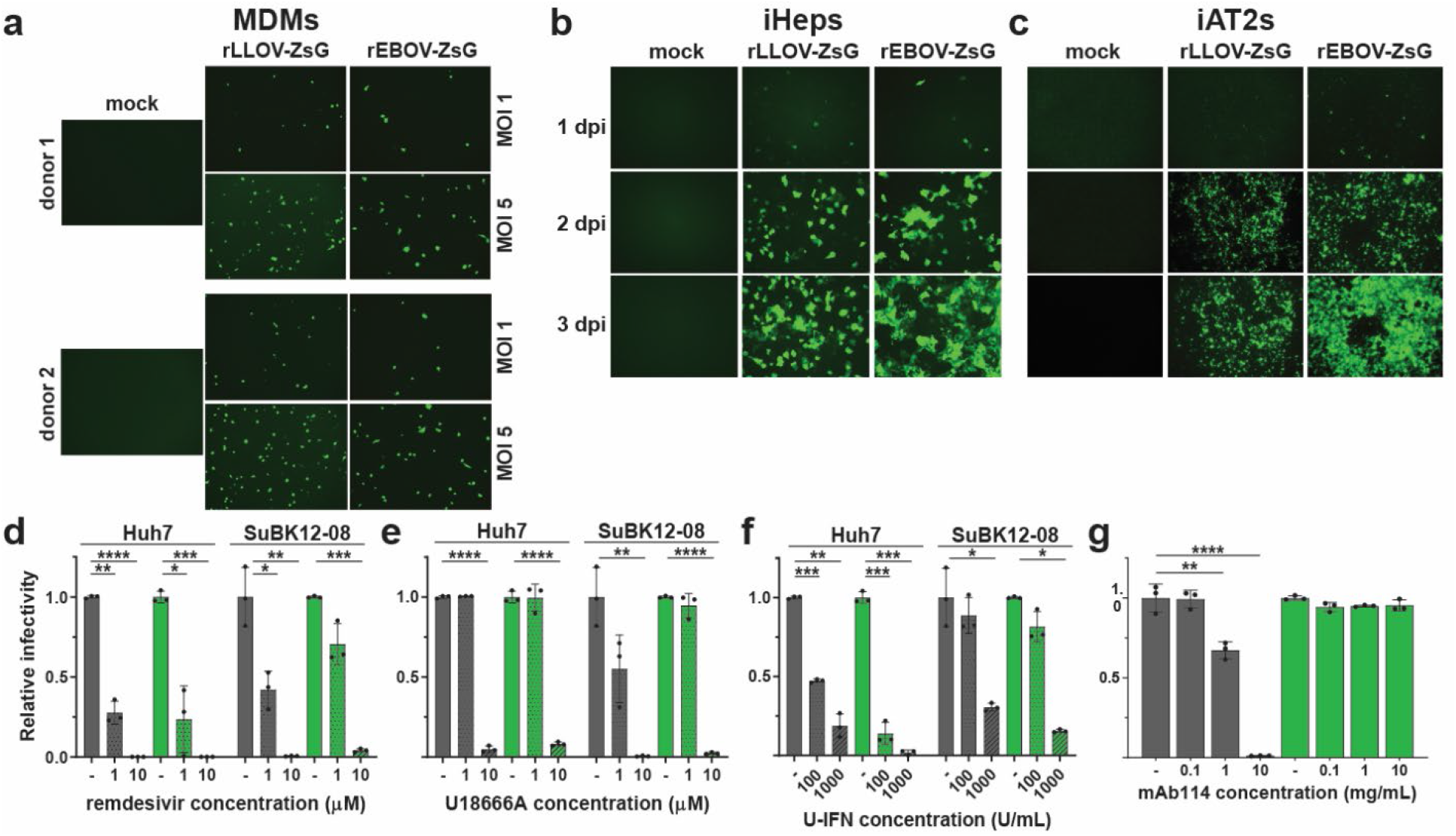
Cellular tropism and antiviral testing of rLLOV_comp_. (**a**) Infection of MDMs derived from 2 donors with the indicated MOIs of rEBOV-ZsG and rLLOV-ZsG-IR_ins_-EBOV_UTR+tr_, labeled rLLOV-ZsG. Fluorescent images taken at 2 dpi. Infections of (**b**) iPSC-derived hepatocytes (iHeps) or (**c**) iPSC-derived lung alveolar type 2 cells (iAT2s) with rEBOV-ZsG or rLLOV-ZsG at an MOI of 1. Fluorescent images taken at the indicated dpi. (**d-f**) Testing antiviral compounds against rEBOV-ZsG (gray bars) and rLLOV-ZsG (green bars) in human and bat cells. Huh7 and SuBK12-08 cells were pre-treated with the indicated concentrations of remdesivir (**d**) or the NPC-1 inhibitor U18666A (**e**) for 30 minutes, or with universal interferon (U-IFN) for 18 hours (**f**) prior to infection with rEBOV-ZsG or rLLOV-ZsG at an MOI of 0.1. Fluorescent images were taken at 2 dpi and mean fluorescence relative to infected cells pre-treated with vehicle control are shown. (**g**) Neutralization assay of rEBOV-ZsG and rLLOV-ZsG at an MOI of 10 using the indicated amounts of EBOV-neutralizing antibody mAb114. Fluorescent images were taken at 2 dpi and relative percentages of infected cells are shown.

Since some of the Schreiber’s bats identified during LLOV-associated die-offs showed lung infiltrates ^2,5^, we also tested the ability of these viruses to infect iPSC-derived human alveolar type 2 cells (iAT2s) ^19,20^, an important lung cell type which had never been previously analyzed in the context of filovirus infections. Both rEBOV and rLLOV_comp_ were able to infect these cells efficiently, with rapid virus spread (Fig. 3c). Although filovirus infections in humans have not been associated with aerosol spread, the lung is a target organ of EBOV infection in humans as shown by pulmonary histopathology of fatal EBOV disease cases ^17^. In addition, EBOV infection through the aerosol route has been demonstrated in nonhuman primates ^21^. EBOV and RESTV infections of pigs were associated with lung pathology and oronasal shedding of viral particles ^22,23^, indicating that there might be host and virus species specific differences in filoviral pathogenesis and transmission routes.

### Use of rLLOV_comp_ to test the efficacy of potential therapeutics

One highly useful application for recombinant viruses such as rLLOV_comp_ that are rescued with the aid of complementation with small non-coding terminal sequences from closely related viruses is the ability to study the effects of antiviral compounds on a highly authentic replication-competent virus. Comparing rLLOV_comp_ and rEBOV in both human and bat cells, we found similar inhibition of each virus in both cell types by remdesivir (Fig. 3d), a nucleoside analog known to inhibit EBOV RNA synthesis ^24^, confirming recently published results using a LLOV minigenome system ^25^. Parallel analysis using U18666A, an inhibitor of the filovirus receptor Niemann-Pick C1 (NPC1) previously shown to inhibit EBOV entry ^26,27^, also showed inhibition of both viruses in both cell types (Fig. 3e), which is in line with previous results indicating that LLOV uses NPC1 to mediate cell entry ^28^. rLLOV_comp_ was slightly more sensitive to universal type I IFN pre-treatment than rEBOV, particularly in human cells (Fig. 3f). Conversely, while mAb114 (brand name Ebanga), a neutralizing monocolonal antibody targeting EBOV GP and approved for treatment of EBOV disease ^29^, effectively blocked rEBOV infection in a dose-dependent manner at concentrations similar to those previously published ^29^, it had no effect on rLLOV_comp_ replication (Fig. 3g). This corroborates previous studies showing that pan-ebolavirus anti-GP antibodies did not block entry of vesicular stomatitis virus pseudotyped with LLOV GP ^30^, emphasizing the requirement of designing antivirals tailored to the virus of interest. In summary, rLLOV_comp_ can be used to test all steps of the viral replication cycle, including entry, viral RNA synthesis, assembly, and egress, as potential targets for antiviral therapeutics using a single system.

### Use of rLLOV_comp_ for host response analysis

A hallmark of severe EBOV infection is an exacerbated inflammatory response characterized by the induction of interferon-stimulated genes and proinflammatory modulators ^8^. This inflammatory response is likely driven by activated monocytes and macrophages ^31^. We previously compared the host response to infection with the highly pathogenic EBOV and the less pathogenic RESTV in human MDMs and observed a strong upregulation of type I IFN and proinflammatory responses in EBOV-infected MDMs, whereas RESTV-infected MDMs remained mainly silent ^32^. To explore whether rLLOV_comp_ would induce an EBOV- or RESTV-like host response in macrophages, MDMs obtained from 3 donors were infected with EBOV, RESTV, or rLLOV_comp_ at an MOI of 10. RNA-FISH analysis demonstrated the presence of viral genome and transcripts for all viruses, yet variable infection rates were observed, with a trend of rLLOV_comp_ having infection rates similar to, or higher than, EBOV and RESTV. However, rLLOV_comp_ infections generally appeared to be in an earlier stage of infection as evidenced by less abundant positive and negative sense viral RNA (Fig. 4a and Supplementary Fig. 6). Infection rates also varied between donors (Fig. 4a). Typical filoviral inclusions within the cytoplasm of the infected MDMs were observed for all viruses as visualized by viral RNA clusters, particularly with viral genomic RNA, indicating robust viral replication in these cells (Fig. 4b).

**Figure 4.**
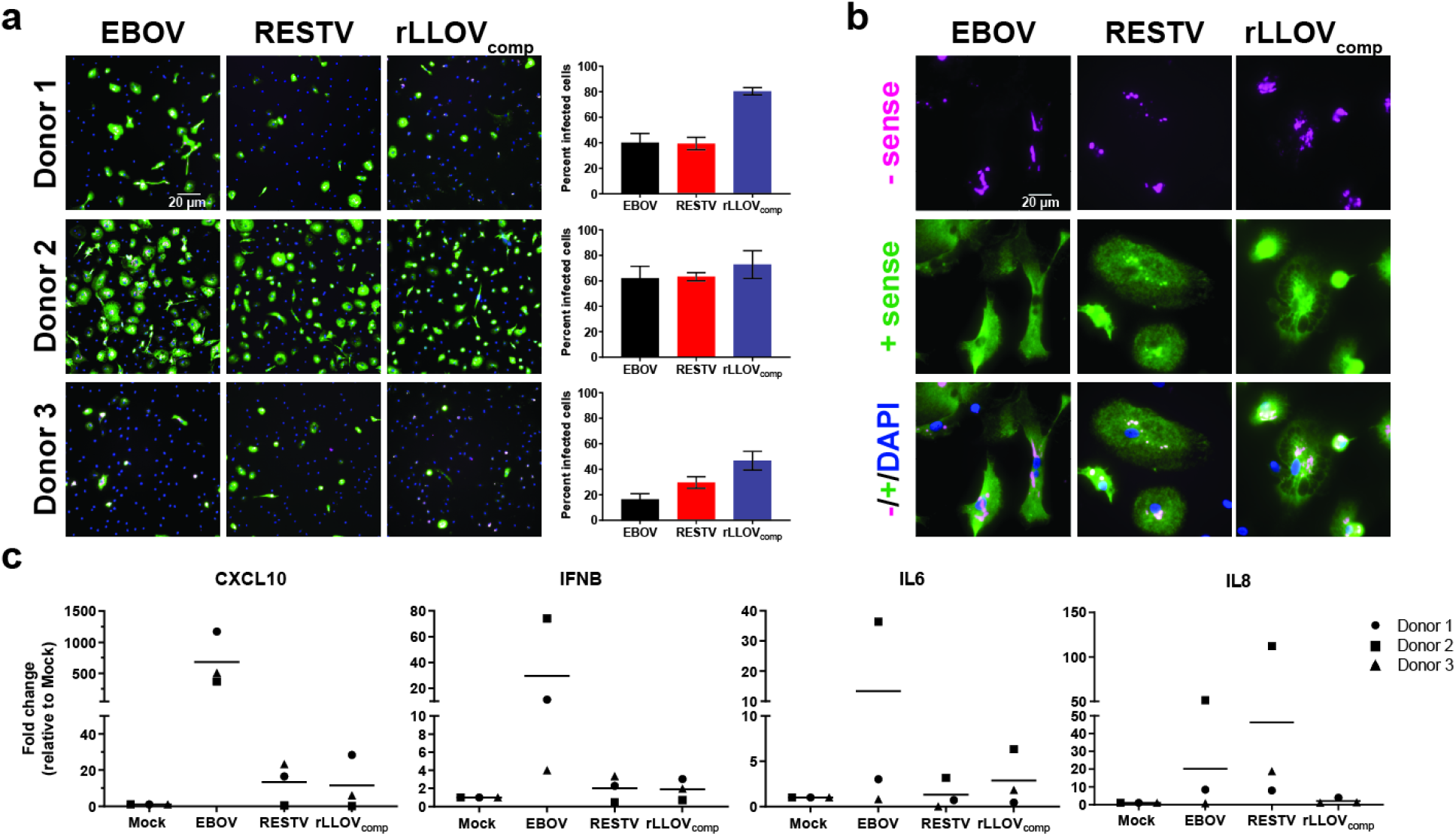
Response to infection with EBOV, RESTV, and rLLOV_comp_ in human monocyte-derived macrophages (MDMs). (**a**) MDMs derived from 3 donors were infected at an MOI of 10 with EBOV, RESTV, or rLLOV-IR_ins_-MARV_UTR+tr_. One dpi, cells were fixed and analyzed for the presence of positive (mRNA and antigenome, green) and negative (genome, magenta) sense viral RNA by RNA FISH. Cells were co-stained with DAPI. 20x images are shown. Quantification of infection rates was determined by RNA FISH, with eight 20x images counted per sample with mean infection rate and SD shown. (**b**) 100x images of the RNA FISH staining in (a) are shown to highlight the association of viral genomic RNA (magenta) with viral inclusions. (**c**) MDMs from the same donors were infected with EBOV, RESTV, and rLLOV-IR_ins_-MARV_UTR+tr_ and total RNA was harvested at 1 dpi. Levels of the indicated analytes were determined by qRT-PCR, with fold change relative to mock-infected cells shown with mean bars.

Transcriptional analysis of select genes confirmed previously observed responses to EBOV and RESTV infection ^32^. While we observed robust upregulation of CXCL10 and IFNβ and, to a lesser extent, upregulation of IL6 and IL8 in EBOV-infected cells, there was only modest or no upregulation of the target genes in RESTV- and rLLOV_comp_-infected cells, suggesting that similar to RESTV, rLLOV_comp_ does not activate human MDMs (Fig. 4c).

Unstudied zoonotic viruses, particularly those closely related to known human pathogens, pose a threat of spillover with the potential to cause epidemics and pandemics within the human population. While many such RNA viruses, including filoviruses, have recently been discovered by unbiased sequencing technologies such as RNA-Seq ^2–4^, the coverage of their genomes is typically incomplete due to missing genomic terminal sequences. Here we provide a blueprint for studying uncultured, potentially pathogenic viruses that currently only exist as incomplete sequences using LLOV, a filovirus, as an example. We used noncoding terminal sequences from other filoviruses to complement the missing LLOV sequence and we also discovered that an intergenic region lacking conserved gene start and gene end signals was nonfunctional. We leveraged the data from these complementation assays to rescue recombinant LLOV. Using rLLOV_comp_, we show that there are no restrictions for LLOV infection of human cells, including primary targets of pathogenic filovirus infections such as macrophages and hepatocytes ^17,33^, seemingly indicating that spillover of LLOV to humans may be possible. While the molecular mechanisms of filovirus pathogenicity still remain elusive, previous studies have hypothesized that slower replication kinetics and muted inflammatory responses with RESTV play a role in the virus being likely nonpathogenic to humans ^32,34,35^. Similarly, LLOV replicates slowly and does not activate MDMs, potentially pointing to LLOV not being pathogenic in humans. Therefore, based on our data with rLLOV_comp_, it appears that LLOV may have a RESTV-like phenotype, having the capacity to infect humans but lacking the ability to cause disease ^36^.

## Discussion

Particularly when trying to answer questions regarding potential pathogenicity, recombinant viruses overcome limitations of other tools that analyze only isolated steps of viral replication, such as viral genome replication and transcription (minigenome assays) or viral entry (virus-like particles) ^37^. Recombinant viruses allow for a plethora of studies to address questions of potential spillover and pathogenicity, including tropism studies, analyzing host responses of primary target cells, and ultimately animal infection studies. Additionally, recombinant viruses are perfectly suited to high throughput drug screens, since they are cheaper, more scalable, and subject to fewer confounding factors such as transfection efficiency that are associated with model systems. The addition of reporter genes and use of RNA-based detection assays are also particularly beneficial for this and other purposes since reagents such as antibodies to detect these novel viruses may not be available.

One limitation to using recombinant viruses with complemented genomic termini is that these exogenous sequences could be suboptimal, making these viruses replicate more slowly and to lower titers. It was therefore surprising that we did not observe substantial sequence changes within the trailer region in the rLLOV_comp_ stocks, suggesting some flexibility of the viral polymerase complex in recognizing promoter elements.

## Materials and Methods

### Biosafety Statement

All work with wildtype and recombinant Ebola virus (EBOV), Reston virus (RESTV) and Lloviu virus (LLOV) was performed in the biosafety level 4 (BSL-4) facility of Boston University’s National Emerging Infectious Diseases Laboratories (NEIDL) following approved standard operating procedures in compliance with local and national regulations pertaining to handling BSL-4 pathogens and Select Agents.

### Cell culture

Cell lines used in this study include human embryonic kidney cells (293T; ATCC CRL-3216, African green monkey kidney cells (Vero E6; ATCC CRL-1586), human liver cells (Huh7; kindly provided by Apath L.L.C.), *Miniopterus schreibersii* kidney cells (SuBK12-08; kindly provided by Ayato Takada, Hokkaido University) ^13^, and golden hamster baby kidney cells (BHK-21; ATCC CCL-10). 293T, Vero E6, and Huh7 cells were maintained in Dulbecco’s modified Eagle medium (DMEM) supplemented with 10% fetal bovine serum (FBS), L-glutamine (200 mM), and either penicillin (50 units/ml) and streptomycin (50 mg/ml) or 100 ug/ml Primocin. BHK-21 cells were maintained in Glasgow’s Minimum Essential Medium (G-MEM) supplemented with 10% FBS, L-glutamine (200 mM), and 2% MEM amino acid solution (50x). SuBK12-08 were grown in Advanced RPMI 1640 supplemented with 10% FBS, 2% MEM amino acid solution (50x), sodium pyruvate (110 mg/L), and 100 ug/ml Primocin. All cell lines were grown at 37°C/5% CO_2_. The following cell lines are female: 293T and Vero E6. The following cell lines are male: BHK-21 and Huh7. The sex of the following cell line is unknown: SuBK12-08.

#### Generation of macrophages

Monocyte-derived macrophages (MDMs) were generated as described previously ^32^. Briefly, MDMs were generated from Leukoreduction system (LRS) chambers (NY Biologics Inc.) using Ficoll separation (GE Healthcare) and isolation of CD14+ monocytes by magnetic bead selection (Milteny Biotech). CD14-selected cells were seeded into ultra-low attachment 6-well plates (Corning) for differentiation into MDMs using RPMI medium supplemented with penicillin (50 units/ml), streptomycin (50 mg/ml), HEPES (10 mM), 5% FBS (certified One Shot, Gibco) supplemented with 100 ng/mL human GMCSF (Peprotech) for 6-8 days. Differentiated MDMs were lifted from plates using Cell Stripper (Corning), and 2×10^5^ cells were seeded into 24-well culture plates or 5×10^4^ cells into Lab-Tek II 8-well chamber slides (Nunc/Thermo Fisher). To account for donor variations, experiments were performed with cells obtained from at least three different donors as noted in the figure legends. Throughout the figures, each donor is represented by a donor-specific symbol.

#### Induced pluripotent stem cell (iPSC) culture and directed differentiation into hepatocytes

The human induced pluripotent stem cell (iPSC) line “BU3” ^38^ was maintained in feeder free conditions on Matrigel (Corning) coated wells in mTeSR-1 media (StemCell Technologies) supplemented with 100 μg/mL Primocin. BU3 was confirmed as mycoplasma free by PCR analysis of gDNA using the following mycoplasma specific primers: 5’ CTT CWT CGA CTT YCA GAC CCA AGG CAT 3’ and 5’ ACA CCA TGG GAG YTG GTA AT 3’ and verified as karyotypically normal as determined by G-band karyotyping analysis from 20 metaphases.

iPSC directed differentiation towards the hepatic lineage was performed using a previously published protocol ^18,39,40^. Briefly, undifferentiated iPSCs were cultured until confluent and then passaged on Day 0 using gentle cell dissociation reagent (GCDR, StemCell Technologies) to achieve single cell suspensions and replated at 1×10^6^ cells per well of a Matrigel coated 6-well plate. Cells were then placed into hypoxic conditions for the remainder of the protocol (5% O_2_, 5% CO_2_, 90% N_2_). Post-passage cells were patterned using the STEMdiff definitive endoderm kit (StemCell Technologies) for 5 days. Day 1 cells were grown in basal media supplemented with STEMdiff™ Definitive Endoderm Supplements MR and CJ followed by an additional 3 days (Days 2-4) in basal media supplemented with CJ alone. At day 5, endoderm was then passaged with GCDR at a 1:4 ratio onto Matrigel-coated wells and cultured in base complete serum-free differentiation medium (cSFDM) supplemented with growth factors as previously described ^18,39,40^. Cells were then cultured for at least an additional 17 days using stage specific growth factors to specify the hepatic lineage and induce maturation.

#### Generation of iAT2 air-liquid interface (ALI) culture

The human iPSC line, SPC2-ST-B2, engineered to carry a tdTomato reporter targeted to the endogenous SFTPC locus ^19^, underwent directed differentiation to generate iPSC-derived alveolar epithelial type 2 cells (iAT2s) in 3D Matrigel cultures using methods we have previously published ^41,42^. Briefly, to establish pure cultures of iAT2s, cells were sorted by flow cytometry to isolate SFTPC^tdTomato+^ cells on day 41 of differentiation, and subsequently maintained through serial passaging as self-renewing monolayered epithelial spheres by plating in Matrigel (Corning) droplets at a density of 400 cells/μl. The cultures were fed every other day with a defined serum-free distal lung maintenance media composed of base cSFDM supplemented with 3 μM CHIR99021, 10 ng/mL KGF, 50 nM dexamethasone, 0.1 mM cAMP, and 0.1 mM IBMX, referred to as “CK+DCI” media and as described previously ^42^. iAT2 culture quality and purity were monitored at each passage by flow cytometry, where > 80% of cells expressing SFTPC^tdTomato^ was observed over time, as shown previously ^19,42^. Air-liquid interface (ALI) cultures were generated as previously described ^20^, with purified iAT2s seeded at 520,000 cells/cm^2^ in 6.5 mm Transwell inserts (Corning) coated with hESC-Qualified Matrigel (Corning) in CK+DCI with 10 μM Rho-associated kinase inhibitor (Sigma Y-27632). At 48h after plating, apical medium is aspirated to initiate air-liquid interface culture, and cultures were fed with CK+DCI basolateral medium every other day prior to experiments.

### Viral sequences

The following NCBI reference filovirus sequences were used for cloning where indicated and for sequence comparison: Lloviu cuevavirus isolate Lloviu virus/M.schreibersii-wt/ESP/2003/Asturias-Bat86 (GenBank: NC_016144), Zaire ebolavirus isolate Ebola virus/H.sapiens-tc/COD/1976/Yambuku-Mayinga (NC_002549), Reston ebolavirus isolate Reston virus/M.fascicularis-tc/USA/1989/Philippines89-Pennsylvania (NC_004161), and Marburg marburgvirus isolate Marburg virus/H.sapiens-tc/KEN/1980/Mt. Elgon-Musoke (NC_001608).

### Plasmids

LLOV support plasmids (pCAGGS-NP_LLOV_, -VP35_LLOV_, -VP30_LLOV_, and -L_LLOV_) and 3L5E-Luc minigenome plasmid were described previously ^9^. EBOV support plasmids (pCAGGS-NP_EBOV_, - VP35_EBOV_, -VP30_EBOV_, and -L_EBOV_) and RESTV support plasmids (pCAGGS-NP_RESTV_, - VP35_RESTV_, -VP30_RESTV_, and -L_RESTV_) were described previously ^9,43^. EBOV support plasmids have been made available through addgene (https://www.addgene.org/Elke_Muhlberger/). A plasmid expressing a codon-optimized version of T7 RNA polymerase was synthesized (GeneArt) and the pMIR β-gal plasmid was a kind gift from Matthew Jones, Boston University.

#### Cloning of for hybrid trailer minigenomes

LLOV minigenomes containing hybrid LLOV L 3’ untranslated region (UTR) and trailer appended with sequences from the EBOV trailer were created as follows. DNA fragments containing the LLOV L 3’ UTR and trailer (nucleotides 18821-18927) appended with the full EBOV L 3’ UTR and trailer (nucleotides 18220-18959) was synthesized (Twist Biosciences) and inserted into the minigenome plasmid 3L5E-Luc ^9^ via NotI and XmaI digestion and ligation (3L5LE_full_-Luc). Truncated versions of this minigenome were generated using the Q5 Site-Directed Mutagenesis Kit (New England Biolabs) using the 3L5LE_full_-Luc as a template.

#### Cloning of bicistronic minigenomes

A LLOV bicistronic minigenome was generated by cloning a synthesized fragment (Twist Biosciences) containing a short fragment of the LLOV 5’ UTR of the nucleoprotein (NP) gene and 3’ UTR of the polymerase (L) gene flanking firefly and renilla luciferase reporters separated by the VP24-L intergenic region (IR) of LLOV (nucleotides 11866-12229, IR_wt_) into minigenome plasmid 3L5LE_72_-Luc using EcoRI and SpeI sites. The IR_ins_ version of the LLOV bicistronic minigenome was generated by replacing the wild-type VP24-L IR region in the LLOV IR_wt_ bicistronic minigenome with a synthesized fragment containing the VP24-L IR with an insertion spanning overlapping LLOV gene start and end signals (GAAGAATATTAAGAAAAA) between nucleotides 138 and 139 of the IR (Twist Biosciences) via NEBuilder HiFi DNA Assembly (New England Biolabs).

#### Cloning of full-length filovirus plasmids

All cloning work with full-length filovirus genomes was performed in the full-length cloning laboratory of the NEIDL. The LLOV full-length clones were generated by inserting the viral genome sequence in plasmid p15AK in positive sense orientation under the control of the T7 RNA polymerase promoter (Supplementary Fig. 1). In this plasmid, the leader region is located immediately downstream of the T7 RNA polymerase promoter and the trailer region is flanked by a hepatitis delta (HdV) ribozyme to generate precise viral genome ends. DNA fragments comprising three portions of the full LLOV genome (NC_016144.1) were synthesized as follows. Fragment 1-LE_72_ (Twist Biosciences) consisted of a NotI site, a T7 RNA polymerase promoter, nucleotides 1-3 of the EBOV genome, nucleotides 1-4202 of the LLOV genome, a linker sequence to facilitate efficient restriction enzyme digestion (CATCGGGCCCCATC), nucleotides 17160-18927 of the LLOV genome including a single silent mutation within the L ORF (A17164T), nucleotides 18887-18958 of the EBOV genome, ending with a portion of the HdV ribozyme SuperCut II (GGGTCGGCATGGCATCTCCACCTCCTCGCGGTCCG). Fragment 2 (Twist Biosciences) consisted of nucleotides 4197-11536 of the LLOV genome. Fragment 3 (Twist Biosciences) consisted of nucleotides 11531-17165 of the LLOV genome containing the same silent mutation within the L ORF as Fragment 1. DNA fragments comprising a portion of the LLOV L open reading frame (nucleotides 18447-18820) followed by EBOV (nucleotides 18220-18958, EBOV_UTR+tr_), or MARV sequences (nucleotides 18477-19111, MARV_UTR+tr_) and ending with same portion of the HdV rizbozyme SuperCutII sequence as used in Fragment 1 were synthesized (Twist Biosciences). A DNA portion of Fragment 2 containing an insertion of the ZsGreen-P2A upstream of the VP40 ORF as described previously ^10^ was synthesized (Twist Biosciences). Finally, a DNA fragment introducing overlapping gene start and end signals into the intergenic region of the VP24-L IR (GAAGAATATTAAGAAAAA between nucleotides 12001 and 12002, IR_ins_) was synthesized.

The LLOV-ZsG-IR_ins_-LE_72_ version full-length clone was assembled by first cloning Fragment 1-LE_72_ into the p15AK-EBOV plasmid (kind gift of Hideki Ebihara, Mayo Clinic) ^44^ via NotI and RsrII digestions and ligation. Fragment 3 was then cloned into p15AK-LLOV-Frag1-LE_72_ by ApaI and AatII digestions and ligation. Finally, Fragment 2 was cloned into p15AK-LLOV-Frag1-3-LE_72_ by XhoI and ApaI digestions and ligation. Other trailer versions of the LLOV full-length clone were assembled by first cloning the LLOV L-EBOV_UTR+tr_ and -MARV_UTR+tr_ fragments into the p15AK-LLOV-Frag1 plasmid via SnaBI and RsrII digestions and ligation. The EBOV_UTR+tr_ and -MARV_UTR+tr_ versions of full-length LLOV clones were assembled by XhoI and AatII digestions and ligation of the backbone portion of the corresponding p15AK-LLOV-Frag1 versions with the Fragment 2+3 portion of the p15AK-LLOV-LE_72_ full length plasmid. ZsGreen-P2A versions of the full-length clone were cloned by first cloning the ZsGreen-P2A fragment into Fragment 2 via XhoI and XmaI digestions and ligation followed by subsequent replacement of this region within the full-length clone via XhoI and ApaI digestions and ligation. IR_ins_ versions of the full-length clone were cloned by first cloning the IR_ins_ fragment into LLOV-Frag3 via ApaI and PciI digestions and ligation followed by subsequent replacement of this region within the full-length clone via ApaI and AatII digestions and ligation.

For the construction of the EBOV-ZsGreen and RESTV-ZsGreen full-length plasmids, DNA fragments consisting of EBOV nucleotides 2148-5773 and RESTV nucleotides 2555-5287 each with an insert of the ZsGreen-P2A upstream of the VP40 ORF as described previously ^10^ were synthesized (Twist Biosciences). These fragments were cloned into the p15AK-EBOV ^44^ and p15AK-RESTV full-length constructs (kind gift of Thomas Hoenen, Friedrich Loeffler Institute) ^35^ via SphI or NcoI and SpeI, respectively.

### Transfections

#### Monocistronic and bicistronic minigenome transfections

LLOV minigenome transfections were performed as described previously ^9^. Briefly, 2×10^5^ 293T or BHK21 cells were seeded in a 12-well plate one day prior to transfection. The next day, cells were transfected with LLOV minigenome plasmid DNA along with LLOV support plasmids encoding NP, VP35, VP30, and L as well as a plasmid expressing codon-optimized T7 RNA polymerase. As negative controls for LLOV minigenome transfections, pCDNA3.1-mCherry plasmid was used instead of L plasmid. When luciferase-expressing minigenomes were used, plasmid pMIR β-gal expressing β-galactose was co-expressed to normalize against transfection efficiency. Transfection of 293T cells was performed using TransIT^®^-LT1 per manufacturer’s recommendations (Mirus Bio LLC).

#### Recombinant filovirus rescue transfections

Recombinant LLOV rescue transfections were performed similar to rescue transfections for other filoviruses ^45^. Briefly, 1:1 mixtures of Huh7:Vero E6 cells (1×10^5^ cells plated per well of a 12-well plate) were transfected with LLOV support plasmids (500 ng pCAGGS-NP_LLOV_, 125 ng pCAGGS-VP35_LLOV_, 50 ng pCAGGS-VP30_LLOV_, and 200 ng pCAGGS-L_LLOV_), pCAGGS-T7 (50 ng; codon-optimized), and a full-length plasmid containing the chimeric LLOV genome (1 μg) using TransIT^®^-LT1 per manufacturer’s recommendations (Mirus Bio LLC). Media was changed approximately 18 hours post-transfection and cells were monitored for cytopathic effect (CPE) and fluorescence for ZsGreen-containing clones. Supernatants of cells showing CPE and/or fluorescence were transferred to T75 flasks of Vero E6 cells approximately 7-11 days post-transfection. rEBOV-ZsGreen and rRESTV-ZsGreen clones were rescued similarly, using the corresponding full-length plasmids (1 μg each), pCAGGS-T7 (50 ng; codon-optimized), plus the EBOV (750 ng pCAGGS-NP_EBOV_, 125 ng pCAGGS-VP35_EBOV_, 50 ng pCAGGS-VP30_EBOV_, and 100 ng pCAGGS-L_EBOV_) or RESTV (350 ng pCAGGS-NP_RESTV_, 125 ng pCAGGS-VP35_RESTV_, 50 ng pCAGGS-VP30_RESTV_, and 350 ng pCAGGS-L_RESTV_) support plasmids, respectively. Rescue of recombinant filoviruses, including LLOV, was performed in BSL-4 facility of the NEIDL following BSL-4 biosafety procedures.

### Viruses

EBOV Mayinga and RESTV Pennsylvania virus isolates were kindly provided by the NIH NIAID Rocky Mountain laboratories. EBOV, RESTV, and LLOV stocks were grown in Vero E6 cells and purified using ultracentrifugation through a 20% sucrose cushion as previously described (*1*). Virus titers were determined in Vero E6 cells by 50% tissue culture infectious dose (TCID_50_) assay and calculated using the Spearman-Kärber algorithm. All work with EBOV, RESTV, and LLOV was performed under BSL-4 conditions at the NEIDL, following approved SOPs.

### Luciferase analysis

#### Monocistronic minigenome firefly luciferase assays

At 3 days post-transfection, cell lysates were harvested in Reporter Lysis Buffer (Promega) and analyzed with the Luciferase Assay System (Promega) using a LUMIstar Omega luminometer (BMG LabTech). Cell lysates were diluted in 1x Reporter Lysis Buffer as needed. To account for potential differences in transfection efficiency, luciferase values were normalized to β-galactosidase values (Promega). Luciferase values were then calculated as fold increase over the negative control (minus L) values.

#### Bicistronic minigenome firefly and renilla luciferase assays

At 3 days post-transfection, cell lysates were harvested in Passive Lysis Buffer (Promega) and analyzed with the Dual-Luciferase Reporter Assay System (Promega) using a LUMIstar Omega luminometer (BMG LabTech). Cell lysates were diluted in 1x Passive Lysis Buffer as needed. To account for potential differences in transfection efficiency, luciferase values were normalized to β-galactosidase values (Promega). Luciferase values were then calculated as fold induction over the negative control (minus L) values.

### Proteomic analysis

#### Sample preparation and mass spectrometry analysis

For proteomic analysis, SuBK12-08 cells seeded in 6-well plates (1.5×10^6^ cells/well) infected with LLOV-ZsG-IR_ins_-EBOV_UTR+tr_ at an MOI of 1 were lysed 2 days post-infection. The virus was inactivated by resuspending the cell pellets in roughly 5 packed cell volumes (p.c.v) of GuHCl lysis buffer (6 M guanidine hydrochloride, 100 mM Tris pH 8.0, 40 mM chloroacetamide, 10 mM Tris(2-carboxyethyl)phosphine) supplemented with phosphatase inhibitor cocktail (Roche) followed by heating to 100°C for 10 minutes. Lysates were removed from the BSL-4 laboratory, sonicated with a Branson probe sonicator and were then quantified via Bradford assay. One hundred sixty micrograms of each sample was diluted with 100 mM Tris buffer, pH 8.5 to decrease the GuHCl concentration to 0.75 M. Lysate proteins were then digested by adding trypsin (Pierce) at a 1:50 ratio (enzyme: protein, w/w) and incubated overnight at 37°C with shaking. Trypsinization was terminated with the addition of trifluoroacetic acid to below pH 3 and the peptide digests were desalted via reversed-phase C18 columns (Sep-pak, Waters) with a wash buffer of 0.1% TFA and elution buffer of 60% acetonitrile. Peptide concentration was determined by a quantitative colorimetric peptide assay (Thermo Fisher), and 1 μg aliquots of clean peptides from each sample were injected sequentially over a nano-flow C18 column (EasySpray, Thermo Fisher) using a EasyNano LC system coupled to an Orbatrap Exploris mass spectrometer (Thermo Fisher) equipped with FAIMS. Data were gathered in a standard data-dependent manner over a 150-minute gradient.

#### Analysis of raw mass spectrometry data

All acquired MS/MS spectra were simultaneously searched against both a manually curated database of LLOV proteins based on the reference sequence (GenBank: NC_016144) and the complete reviewed and unreviewed proteome for the common bats family (uniprotKB taxonomy Vespertilionidae [9431], downloaded February 9, 2021) using the MaxQuant software (Version 1.6.7.0) that integrates the Andromeda search engine. Briefly, enzyme specificity was set to trypsin and up to two missed cleavages were allowed. Cysteine carbamidomethylation was specified as fixed modification whereas oxidation of methionine and N-terminal protein acetylation were set as variable modifications. Precursor ions were searched with a maximum mass deviation of 4.5 ppm and fragment ions with a maximum mass deviation of 20 ppm. Peptide and protein identifications were filtered at 1% FDR using the target-decoy database search strategy ^46^. Proteins that could not be differentiated based on MS/MS spectra alone were grouped to protein groups (default MaxQuant settings). A threshold Andromeda score of 40 was applied to peptides. The MaxQuant output file designated “proteinGroups” was used for data normalization.

### Electron microscopy/cryoelectron microscopy

#### Transmission electron microscopy of filovirus-infected cells

Briefly, Huh7 and SuBK12-08 cells seeded on coverslips in 6-well plates were mock-infected or infected with EBOV or LLOV-IR_ins_-MARV_UTR+tr_ at an MOI of 5. At 1, 2, and 3 dpi, cells were fixed in Karnovsky’s fixative (Electron Microscopy Sciences, EMS) for 6 hours following approved inactivation protocols and removed from the BSL-4 facility. The fixed cells were washed 3 times in 0.1 M cacodylate buffer, postfixed in 1% osmium tetroxide (OsO4)/1.5% potassium ferrocyanide (K_4_[Fe(CN)_6_]) for 30 minutes, washed twice in water, incubated in 1% tannic acid in water for 30 minutes, washed twice in water and once in 50 mM maleate buffer pH 5.15 (MB) followed by a 30-minute incubation in 1% uranyl acetate in MB. The cells were then washed in MB and water and subsequently dehydrated in grades of alcohol (5 minutes each; 50%, 70%, 95%, 2× 100%). Cells were embedded in plastic by inverting a gelatin capsule filled with Epon/Araldite on top of the coverslip and polymerized at 60°C for 24 hours. After polymerization, the coverslip was removed by dipping the block in LN_2_.

Ultrathin sections (about 80 nm) were cut on a Reichert Ultracut-S microtome, picked up onto copper grids, stained with lead citrate and examined in a JEOL 1200EX Transmission electron microscope. Images were recorded with an AMT 2k CCD camera.

#### Transmission electron microscopy and cryoelectron microscopy of filovirus particles

Electron microscopy (EM) and cryo-EM of virion particles were performed as follows. Sucrose cushion-purified stocks of EBOV or LLOV-IR_ins_-MARV_UTR+tr_ were fixed by adding an equal volume of 20% formalin (Fisher Scientific) for at least 6 hours according to approved inactivation procedures. For negatively stained samples, formalin-fixed EBOV or LLOV (4 μl) was applied to the carbon-coated side of the grid and incubated for 3-5 minutes at room temperature. The grid containing virions was then washed on a 0.1 mL drop of Tris buffer (50 mM Tris, pH 7.6) and blotted briefly using filter paper, two times. Grids were then stained on a 5 μl drop of 1% uranyl acetate, blotted briefly, stained again, blotted, and air dried. For cryo-EM, formalin-fixed Ebola or Lloviu virions (1.8 μl) were applied to Quantifoil 2/2 grids, blotted for 7 seconds, and plunge-frozen in liquid ethane using a Vitrobot Mark III. Transmission EM and cryo-EM were performed on an FEI TF20 EM operated at 160 kV and images were recorded on a TVIPS F416 CMOS camera with a pixel size of 1.68 - 10.09 Å, using the software SerialEM.

### Antiviral drug testing

For small molecule testing, 2×10^4^ cells/well of Huh7 and SuBK12-08 cells seeded in 96-well plates were pretreated with the indicated amounts of universal interferon (U-IFN, PBL Assay Science) for 18 hours or remdesivir (Selleck Chemicals Llc), U18666A (Sigma-Aldrich), or DMSO for 30 minutes prior to infection. The cells were then infected with either EBOV-ZsGreen or LLOV-ZsG-IR_ins_-EBOV_UTR+tr_ at an MOI of 1. Fluorescent and brightfield images of infected cells were taken at 2 days post infection. Quantification of fluorescence was performed by subtracting background fluorescence (average intensity of uninfected cells) from the average fluorescence intensity of individual wells. Samples were performed in triplicate.

Testing of mAb114 ^29,47,48^ (brand name Ebanga; kindly provided by Nancy Sullivan, NIH NIAID Vaccine Research Center) was performed by pre-treating 2×10^5^ TCID_50_ units of EBOV-ZsGreen or LLOV-ZsG-IR_ins_-EBOV_UTR+tr_ with media alone or media containing 200 ng/mL, 2 μg/mL, or 20 μg/mL mAb114 at 37° for 1 hour. Following pre-incubation, cells were infected with these mixtures, followed by removal of the inocula after 1 hour and replacement with fresh media. Cells were fixed at 2 dpi in 10% formalin (Fisher Scientific) for at least 6 hours. Samples were stained with 4’,6-diamidino-2-phenylindole (DAPI; 200 ng/mL and images covering at least 90% of each well’s area at 4x magnification were acquired using a Nikon Ti2 Eclipse microscope and Photometrics Prime BSI camera with NIS Elements AR software. Images were stitched, then imported into QuPath software ^49^. Infection rates were determined using the positive cell detection feature in QuPath. Nuclei were detected in the DAPI channel and infected cells were identified by mean fluorescence value in the cytoplasmic region in the green channel.

### RNA fluorescence in situ hybridization analysis

For RNA fluorescent in situ hybridization (FISH) analysis, MDMs seeded in 8-well chamber slides were mock-infected or infected with the indicated viruses at an MOI of 10. Cells were fixed one day post-infection in 10% formalin for at least 6 hours. RNA FISH was performed using the RNAscope Multiplex Fluorescent V2 kit (Advanced Cell Diagnostics). Filovirus RNA was detected using custom-designed probes (Advanced Cell Diagnostics). Specifically, viral mRNA of each filovirus was detected in channel 1 using probes targeting the VP35 transcripts (Advanced Cell Diagnostics) and stained with Opal 520 fluorophore (Perkin-Elmer). Viral genomic RNA was detected in channel 3 using probes targeting the negative-sense genomic sequences of the NP gene (Advanced Cell Diagnostics) and stained with Opal 690 fluorophore (Perkin-Elmer). Staining was performed according to the manufacturer’s protocol for adherent cell samples, with the exception of an additional HRP blocking step following the signal development of the probes detecting viral mRNA as per the manufacturer’s recommendation. Nuclei were stained with kit-supplied DAPI following the manufacturer’s protocol. Coverslips were mounted on slides using FluorSave mounting medium, and slides were subsequently stored at 4°C prior to imaging. Images were acquired at 20x magnification from sixteen fields of view per well in two replicate wells of each chamber slide using a Nikon Ti2 Eclipse microscope and Photometrics Prime BSI camera with NIS Elements AR software. Total cell number was determined from images using the QuPath cell detection feature to identify nuclei by DAPI staining, and infected cells were manually identified based on the presence of staining for viral mRNA or genomic RNA.

### RNA sample preparation

Total RNA from 2×10^5^ (24-well format) MDMs was isolated at one day post-infection using TRIzol reagent (Invitrogen) according to the manufacturer’s protocol. RNA concentration was determined by measuring light absorption at a wavelength of 260 nm using a NanoDrop 1000 spectrophotometer (Thermo Fisher).

### qRT-PCR analysis

Five nanograms of mRNA were analyzed by quantitative reverse transcription-PCR (qRT-PCR) using the QuantiFast SYBR green RT-PCR kit (Qiagen). qRT-PCR-based quantification of IL-6, IL-8, IFNB, CXCL10, and β2-microglobulin (B2M) mRNAs were performed using validated QuantiTect primer assays (Qiagen). B2M levels were used as an endogenous control for normalization. qRT-PCR was performed in a CFX96 real-time PCR cycler (Bio-Rad). Fold expression levels compared to those of non-infected control samples were quantified using the threshold cycle (ΔΔCT) method (Bio-Rad CFX Manager 1.5 software) for each donor.

### Viral RNA sequencing

For viral RNA sequencing, total RNA from sucrose cushion-purified viral stocks was isolated using TRIzol-LS reagent (Invitrogen) according to the manufacturer’s protocol. RNA concentration was determined using a NanoDrop 1000 spectrophotometer (Thermo Fisher).

#### RNA-Seq

Sequencing libraries were prepared using the Illumina TruSeq v2 library kit without rRNA depletion or mRNA selection and were sequenced on the Illumina HiSeq 500 to a depth of > 500,000 reads per library. Raw FASTQ files were evaluated for quality using FastQC v0.11.7 ^50^ followed by alignment to the Vero E6 genome (ChlSab1.1a) using Bowtie2 v2.4.2 ^51^ with the very-sensitive flag. Reads that did not align to the host genome were passed to Kraken2 v2.0.9 ^52^ to verify no adventitious agents or other contaminants were present; none were detected. The reads were then aligned to the predicted viral genome using Bowtie2. Viral coverage was assessed with SAMtools v1.10 ^53^, and single nucleotide variants (SNVs) were called with LoFreq v2.1.3.1 ^54^. SNV annotation was performed with Biostrings v2.56.0 ^55^ and a custom R script.

#### 5’-end genomic sequencing with ViBE-Seq

Viral genomic end sequencing was performed using ViBE-Seq (sequencing of <u>Vi</u>ral *<u>B</u>onafide* <u>E</u>nds), which is a novel method based on the adaptor ligation and cDNA recircularization approaches described previously ^56–58^. Briefly, 40 ng of RNA was used as a template for reverse transcription with QIAGEN’s Sensiscript Reverse Transcription Kit according to the manufacturer’s instructions. Specific RT primers were used for cDNA synthesis for each rLLOV-ZsG-IR_ins_-EBOV_UTR+tr_ (/5Phos/T*A*TGTGAGTCGTATTACCTG/iSp18/GGATCC/iSp18/CCT TCTTTACAATATAGCAGA) and rLLOV-ZsG-IR_ins_-MARV_UTR+tr_ (/5Phos/T*A*TGTGAGTCG TATTACCTG/iSp18/GGATCC/iSp18/GCTGTAATCTACAAGCACCTCTT), where * denotes a non-scissile phosphorothioate linkage and iSp18 is an 18-atom hexa-ethyleneglycol spacer. Reverse transcription products were run on a 10% denaturing urea polyacrylamide gel for 2 hours at 10 watts. The gel was then stained with SYBR Gold for 10 minutes and gel portions corresponding to 200 nucleotides (rLLOV-ZsG-IR_ins_-EBOV_UTR+tr_) or 230 nucleotides (rLLOV-ZsG-IR_ins_-MARV_UTR+tr_) were excised. The bottom of a 0.2 mL tube was punctured with a 21 gauge needle. Excised gel bands were extruded via centrifugation through the punctured 0.2 mL tube at 21,000 *x g* for 5 minutes. The pulverized polyacrylamide was resuspended in 0.5 mL of 0.4 M NaCl solution and shaken overnight at 4°C. Polyacrylamide was eliminated from the slurry via centrifugation through a Spin-X centrifuge tube filter (Costar). cDNA was then precipitated by adding 1 μL of Glycoblue Coprecipitant (Thermo Fisher) and 1 volume of 100% isopropanol, overnight incubation at −20°C, and centrifugation at 21,000 *x g* for 1 hour at 4°C. The pellet was washed with 1 mL of 100% ethanol, allowed to air-dry, and resuspended in 20 μL of nuclease-free water. cDNA was quantified using a NanoDrop 1000 spectrophotometer (Thermo Fisher).

Purified cDNA was subject to circularization via CircLigase ssDNA Ligase (Lucigen) per the manufacturer’s specifications. Samples were incubated for 2 hours at 60°C and 10 minutes at 80°C.

PCR reactions to amplify the trailer sequences from the circularized cDNA using Q5 High-Fidelity DNA Polymerase (New England Biolabs) using 60 ng of circularized template were then set up with the following parameters: 98°C (1 min), [98°C (10s), 43°C (30s), 72°C (12s), repeat 3-5x,] [98°C (10s), 55°C (30s), 72°C (12s), repeat 29x,] 72°C (2 min), 4°C (hold). For each virus, an extended T7 complement forward primer (GCGTCAGTCAGGTAATACGACTCACATA) was used in combination with a virus-specific reverse primer (TACCTTCTTTACAATATAGCAGACTAGATAATAATCTTCGTGTT for rLLOV-ZsG-IR_ins_-EBOV_UTR+tr_ and GCTGTAATCTACAAGCACCTCTTTTAAATACATTAGGAA for rLLOV-ZsG-IR_ins_-MARV_UTR+tr_). PCR products were run on a 2% agarose gel at 100 V for 1 hour. Bands at 200 nucleotides (rLLOV-ZsG-IR_ins_-EBOV_UTR+tr_) or 230 nucleotides (rLLOV-ZsG-IR_ins_-MARV_UTR+tr_) were excised and purified via Monarch DNA Gel Extraction Kit (New England Biolabs) according to the manufacturer’s instructions. PCR reactions were repeated as necessary to obtain 5-10 ng/μl in 20 μL of nuclease-free water.

Purified PCR products were submitted for sequencing at Mass General Hospital’s Center for Computation and Integrative Biology DNA Core in Cambridge, MA.

### Statistics

Luciferase assays and drug testing: Standard error of the mean (SEM) and two-tailed *t-*tests for all figures were calculated using GraphPad Prism software.

## Acknowledgments

The authors wish to thank Thomas Hoenen and Hideki Ebihara for sharing recombinant wildtype EBOV and RESTV cDNA clones, Nancy Sullivan for donation of antibody mAB114, Ayato Takada for sharing SuBK12-08 cells, Apath L.L.C. for sharing Huh7 cells, Matthew Jones for sharing plasmid pMIR-β-gal, and the National Institutes of Health (NIH) Rocky Mountain Laboratories for providing EBOV and RESTV stocks. This work was supported by NIH grants R21 AI137793 (EM), R01 AI133486 (EM), R21 AI135912 (EM), R21 AI147285 (DC and EM), R01 GM102474 (EB), R01HL095993 (DNK), U01TR001810 (AAW), R01DK101501 (AAW), and R01DK117940 (AAW). iPSC distribution and disease modeling was supported by NIH grants U01TR001810, and N01 75N92020C00005. ELS was supported by NIH training grant T32 HL007035, and GK was supported by the Janos Bolyai Research Scholarship of the Hungarian Academy of Sciences.

## Author contributions

AJH and BH performed most of the experiments. BSL-4 experiments were performed by AJH with help from JO and BH, minigenome experiments were performed by AJH, BH, and WAS, RNA FISH analysis was performed by ELS, quantification of infected cells was performed by ELS and AJH, mass spectrometry and proteomic analysis was performed by RMH, electron microscopy analysis was performed by EB, ME, and MRW, RT-PCR analysis was performed by BH and JO, MDM differentiation was performed by JO and MRW, iAT2 generation was performed by JH and KDA, iHep generation was performed by JEK, RNA-Seq was performed by JT, viral genomic end sequencing was performed by SR, cloning of EBOV-ZsGreen was performed by TTNT, cloning of all other full-length filovirus constructs was performed by BH and AJH. GET, GK, and FJ provided LLOV RNA sequence information. DC, JHC, AE, AAW, DNK, VT, HF, and GK assisted with experimental planning, data analysis and interpretation. AJH and EM conceived the study. The manuscript was written by AJH and EM with assistance from the coauthors.

## Materials & Correspondence

Correspondence and material requests should be addressed to EM.

## References

1 Letko, M., Seifert, S. N., Olival, K. J., Plowright, R. K. & Munster, V. J. Bat-borne virus diversity, spillover and emergence. Nat Rev Microbiol 18, 461–471, doi:10.1038/s41579-020-0394-z (2020).

2 Negredo, A. et al. Discovery of an ebolavirus-like filovirus in europe. PLoS Pathog 7, e1002304, doi:10.1371/journal.ppat.1002304 PPATHOGENS-D-11-01432 [pii] (2011).

3 Goldstein, T. et al. The discovery of Bombali virus adds further support for bats as hosts of ebolaviruses. Nat Microbiol 3, 1084–1089, doi:10.1038/s41564-018-0227-2 (2018).

4 Yang, X. L. et al. Characterization of a filovirus (Mengla virus) from Rousettus bats in China. Nat Microbiol, doi:10.1038/s41564-018-0328-y (2019).

5 Kemenesi, G. et al. Re-emergence of Lloviu virus in Miniopterus schreibersii bats, Hungary, 2016. Emerg Microbes Infect 7, 66, doi:10.1038/s41426-018-0067-4 (2018).

6 Ramirez de Arellano, E. et al. First Evidence of Antibodies Against Lloviu Virus in Schreiber’s Bent-Winged Insectivorous Bats Demonstrate a Wide Circulation of the Virus in Spain. Viruses 11, doi:10.3390/v11040360 (2019).

7 Amarasinghe, G. K. et al. Taxonomy of the order Mononegavirales: update 2019. Arch Virol 164, 1967–1980, doi:10.1007/s00705-019-04247-4 (2019).

8 Liu, X. et al. Transcriptomic signatures differentiate survival from fatal outcomes in humans infected with Ebola virus. Genome Biol 18, 4, doi:10.1186/s13059-016-1137-3 (2017).

9 Manhart, W. A. et al. A Chimeric Lloviu Virus Minigenome System Reveals that the Bat-Derived Filovirus Replicates More Similarly to Ebolaviruses than Marburgviruses. Cell Rep 24, 2573–2580 e2574, doi:10.1016/j.celrep.2018.08.008 (2018).

10 Albarino, C. G., Wiggleton Guerrero, L., Lo, M. K., Nichol, S. T. & Towner, J. S. Development of a reverse genetics system to generate a recombinant Ebola virus Makona expressing a green fluorescent protein. Virology 484, 259–264, doi:10.1016/j.virol.2015.06.013 (2015).

11 Whelan, S. P., Barr, J. N. & Wertz, G. W. Transcription and replication of nonsegmented negative-strand RNA viruses. Curr Top Microbiol Immunol 283, 61–119 (2004).

12 Schneemann, A., Schneider, P. A., Lamb, R. A. & Lipkin, W. I. The remarkable coding strategy of borna disease virus: a new member of the nonsegmented negative strand RNA viruses. Virology 210, 1–8, doi:10.1006/viro.1995.1311 (1995).

13 Takadate, Y. et al. Receptor-Mediated Host Cell Preference of a Bat-Derived Filovirus, Lloviu Virus. Microorganisms 8, doi:10.3390/microorganisms8101530 (2020).

14 Shabman, R. S. et al. Deep sequencing identifies noncanonical editing of Ebola and Marburg virus RNAs in infected cells. MBio 5, e02011, doi:10.1128/mBio.02011-14 (2014).

15 Kolesnikova, L., Nanbo, A., Becker, S. & Kawaoka, Y. Inside the Cell: Assembly of Filoviruses. Curr Top Microbiol Immunol 411, 353–380, doi:10.1007/82_2017_15 (2017).

16 Messaoudi, I., Amarasinghe, G. K. & Basler, C. F. Filovirus pathogenesis and immune evasion: insights from Ebola virus and Marburg virus. Nat Rev Microbiol 13, 663–676, doi:10.1038/nrmicro3524 (2015).

17 Martines, R. B., Ng, D. L., Greer, P. W., Rollin, P. E. & Zaki, S. R. Tissue and cellular tropism, pathology and pathogenesis of Ebola and Marburg viruses. J Pathol 235, 153–174, doi: 10.1002/path.4456 (2015).

18 Kaserman, J. E. et al. A Highly Phenotyped Open Access Repository of Alpha-1 Antitrypsin Deficiency Pluripotent Stem Cells. Stem Cell Reports 15, 242–255, doi:10.1016/j.stemcr.2020.06.006 (2020).

19 Hurley, K. et al. Reconstructed Single-Cell Fate Trajectories Define Lineage Plasticity Windows during Differentiation of Human PSC-Derived Distal Lung Progenitors. Cell Stem Cell 26, 593–608 e598, doi:10.1016/j.stem.2019.12.009 (2020).

20 Huang, J. et al. SARS-CoV-2 Infection of Pluripotent Stem Cell-Derived Human Lung Alveolar Type 2 Cells Elicits a Rapid Epithelial-Intrinsic Inflammatory Response. Cell Stem Cell, doi:10.1016/j.stem.2020.09.013 (2020).

21 Reed, D. S., Lackemeyer, M. G., Garza, N. L., Sullivan, L. J. & Nichols, D. K. Aerosol exposure to Zaire ebolavirus in three nonhuman primate species: differences in disease course and clinical pathology. Microbes Infect 13, 930–936, doi:10.1016/j.micinf.2011.05.002 (2011).

22 Kobinger, G. P. et al. Replication, pathogenicity, shedding, and transmission of Zaire ebolavirus in pigs. J Infect Dis 204, 200–208, doi:10.1093/infdis/jir077 (2011).

23 Haddock, E. et al. Reston virus causes severe respiratory disease in young domestic pigs. Proc Natl Acad Sci U S A 118, doi:10.1073/pnas.2015657118 (2021).

24 Warren, T. K. et al. Therapeutic efficacy of the small molecule GS-5734 against Ebola virus in rhesus monkeys. Nature 531, 381–385, doi:10.1038/nature17180 (2016).

25 Bodmer, B. S. et al. Remdesivir inhibits the polymerases of the novel filoviruses Lloviu and Bombali virus. Antiviral Res 192, 105120, doi:10.1016/j.antiviral.2021.105120 (2021).

26 Carette, J. E. et al. Ebola virus entry requires the cholesterol transporter Niemann-Pick C1. Nature 477, 340–343, doi:10.1038/nature1034 nature10348 [pii] (2011).

27 Shoemaker, C. J. et al. Multiple cationic amphiphiles induce a Niemann-Pick C phenotype and inhibit Ebola virus entry and infection. PLoS One 8, e56265, doi:10.1371/journal.pone.005626 PONE-D-12-34225 [pii] (2013).

28 Ng, M. et al. Cell entry by a novel European filovirus requires host endosomal cysteine proteases and Niemann-Pick C1. Virology 468-470, 637–646, doi:10.1016/j.virol.2014.08.019 (2014).

29 Misasi, J. et al. Structural and molecular basis for Ebola virus neutralization by protective human antibodies. Science 351, 1343–1346, doi:10.1126/science.aad6117 (2016).

30 Wec, A. Z. et al. Antibodies from a Human Survivor Define Sites of Vulnerability for Broad Protection against Ebolaviruses. Cell 169, 878–890 e815, doi:10.1016/j.cell.2017.04.037 (2017).

31 Kotliar, D. et al. Single-Cell Profiling of Ebola Virus Disease In Vivo Reveals Viral and Host Dynamics. Cell, doi:10.1016/j.cell.2020.10.002 (2020).

32 Olejnik, J. et al. Ebolaviruses Associated with Differential Pathogenicity Induce Distinct Host Responses in Human Macrophages. J Virol 91, doi:10.1128/JVI.00179-17 (2017).

33 Zaki, S. R. & Goldsmith, C. S. Pathologic features of filovirus infections in humans. Curr Top Microbiol Immunol 235, 97–116 (1999).

34 Bodmer, B. S. et al. Differences in Viral RNA Synthesis but Not Budding or Entry Contribute to the In Vitro Attenuation of Reston Virus Compared to Ebola Virus. Microorganisms 8, doi:10.3390/microorganisms8081215 (2020).

35 Groseth, A. et al. The Ebola virus glycoprotein contributes to but is not sufficient for virulence in vivo. PLoS Pathog 8, e1002847, doi:10.1371/journal.ppat.1002847 (2012).

36 Miranda, M. E. & Miranda, N. L. Reston ebolavirus in humans and animals in the Philippines: a review. J Infect Dis 204 Suppl 3, S757–760, doi:10.1093/infdis/jir296 (2011).

37 Maruyama, J. et al. Characterization of the envelope glycoprotein of a novel filovirus, lloviu virus. J Virol 88, 99–109, doi:10.1128/JVI.02265-13 (2014).

38 Kurmann, A. A. et al. Regeneration of Thyroid Function by Transplantation of Differentiated Pluripotent Stem Cells. Cell Stem Cell 17, 527–542, doi:10.1016/j.stem.2015.09.004 (2015).

39 Kaserman, J. E. & Wilson, A. A. Protocol for Directed Differentiation of Human Induced Pluripotent Stem Cells (iPSCs) to a Hepatic Lineage. Methods Mol Biol 1639, 151–160, doi:10.1007/978-1-4939-7163-3_15 (2017).

40 Wilson, A. A. et al. Emergence of a stage-dependent human liver disease signature with directed differentiation of alpha-1 antitrypsin-deficient iPS cells. Stem Cell Reports 4, 873–885, doi:10.1016/j.stemcr.2015.02.021 (2015).

41 Jacob, A. et al. Differentiation of Human Pluripotent Stem Cells into Functional Lung Alveolar Epithelial Cells. Cell Stem Cell 21, 472–488 e410, doi:10.1016/j.stem.2017.08.014 (2017).

42 Jacob, A. et al. Derivation of self-renewing lung alveolar epithelial type II cells from human pluripotent stem cells. Nat Protoc 14, 3303–3332, doi:10.1038/s41596-019-0220-0 (2019).

43 Schümann, M., Gantke, T. & Mühlberger, E. Ebola virus VP35 antagonizes PKR activity through its C-terminal interferon inhibitory domain. J Virol 83, 8993–8997, doi:JVI.00523-09 [pii] 10.1128/JVI.00523-09 (2009).

44 Tsuda, Y. et al. An Improved Reverse Genetics System to Overcome Cell-Type-Dependent Ebola Virus Genome Plasticity. J Infect Dis 212 Suppl 2, S129–137, doi:10.1093/infdis/jiu681 (2015).

45 Brauburger, K. et al. Analysis of the highly diverse gene borders in Ebola virus reveals a distinct mechanism of transcriptional regulation. J Virol 88, 12558–12571, doi:10.1128/JVI.01863-14 (2014).

46 Elias, J. E. & Gygi, S. P. Target-decoy search strategy for increased confidence in large-scale protein identifications by mass spectrometry. Nat Methods 4, 207–214, doi:10.1038/nmeth1019 (2007).

47 Corti, D. et al. Protective monotherapy against lethal Ebola virus infection by a potently neutralizing antibody. Science 351, 1339–1342, doi:10.1126/science.aad5224 (2016).

48 Lee, A. Ansuvimab: First Approval. Drugs 81, 595–598, doi:10.1007/s40265-021-01483-4 (2021).

49 Bankhead, P. et al. QuPath: Open source software for digital pathology image analysis. Sci Rep 7, 16878, doi:10.1038/s41598-017-17204-5 (2017).

50 Andrews, S. FastQC: A Quality Control Tool for High Throughput Sequencing Data. (2015).

51 Langmead, B. & Salzberg, S. L. Fast gapped-read alignment with Bowtie 2. Nat Methods 9, 357–359, doi:10.1038/nmeth.1923 (2012).

52 Wood, D. E., Lu, J. & Langmead, B. Improved metagenomic analysis with Kraken 2. Genome Biol 20, 257, doi:10.1186/s13059-019-1891-0 (2019).

53 Li, H. et al. The Sequence Alignment/Map format and SAMtools. Bioinformatics 25, 2078–2079, doi: 10.1093/bioinformatics/btp352 (2009).

54 Wilm, A. et al. LoFreq: a sequence-quality aware, ultra-sensitive variant caller for uncovering cell-population heterogeneity from high-throughput sequencing datasets. Nucleic Acids Res 40, 11189–11201, doi:10.1093/nar/gks918 (2012).

55 Pagès, H., Aboyoun, R., Gentleman, R. & DebRoy, S. Biostrings: Efficient manipulation of biological strings. R package version 2.56.0 (2020).

56 Huppertz, I. et al. iCLIP: protein-RNA interactions at nucleotide resolution. Methods 65, 274–287, doi:10.1016/j.ymeth.2013.10.011 (2014).

57 McGlincy, N. J. & Ingolia, N. T. Transcriptome-wide measurement of translation by ribosome profiling. Methods 126, 112–129, doi:10.1016/j.ymeth.2017.05.028 (2017).

58 Kretov, D. A. et al. Ago2-Dependent Processing Allows miR-451 to Evade the Global MicroRNA Turnover Elicited during Erythropoiesis. Mol Cell 78, 317–328 e316, doi:10.1016/j.molcel.2020.02.020 (2020).

59 Ellis, D. S. et al. Ebola and Marburg viruses: I. Some ultrastructural differences between strains when grown in Vero cells. J Med Virol 4, 201–211 (1979).

