## Supplementary Data for "Recombinant Lloviu virus as a model to study inaccessible zoonotic viruses"

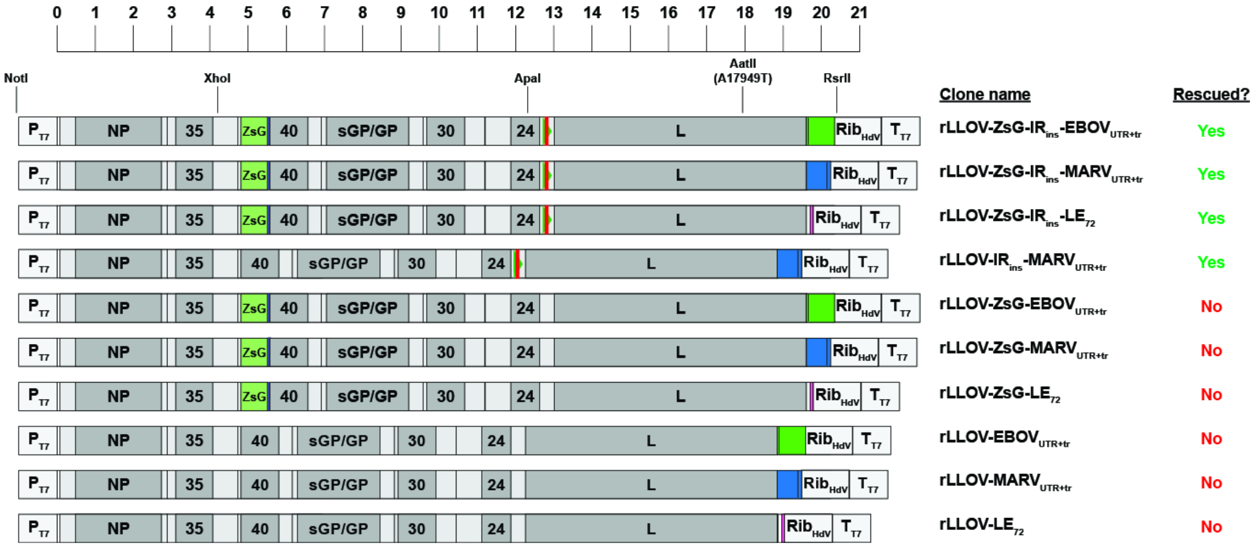


### Supplementary Figure 1

**Recombinant Lloviu virus (rLLOV$_{comp}$) clones used in this study.** Schematics and names of full-length clones of rLLOV$_{comp}$ that were rescued (above) and those that were not (below). Noncoding regions are indicated in light gray, LLOV genes are in gray, a ZsGreen-P2A reporter (ZsG) is in light green followed by a dark gray bar, gene start and end signals are black bars within noncoding regions, red bars indicate the IR$_{ins}$ insertion in the VP24-L intergenic region, dark green indicates the Ebola virus (EBOV) L 3' UTR and trailer (EBOV$_{UTR+tr}$), dark blue indicates the Marburg virus L 3' UTR and trailer (MARV$_{UTR+tr}$), and pink indicates the last 72 nucleotides of the EBOV trailer (LE$_{72}$). rLLOV$_{comp}$ clones are to scale with the exception of the T7 RNA polymerase promoter (P$_{T7}$), hepatitis delta virus ribozyme (Rib$_{HdV}$), and T7 RNA polymerase terminator sequences (T$_{T7}$) which are enlarged for clarity. Scale bar above indicates 1 kb increments, starting with the first nucleotide of the rLLOV$_{comp}$ clones. Important restriction enzymes that were used to clone full-length rLLOV$_{comp}$ plasmids are indicated above, including an AatII site which introduces a silent mutation within L (A17949T for the rLLOV-ZsG-IR$_{ins}$-EBOV$_{UTR+tr}$ clone, corresponding to nucleotide 17164 of the published LLOV sequence, NCBI Reference Sequence NC_016144).

**a**

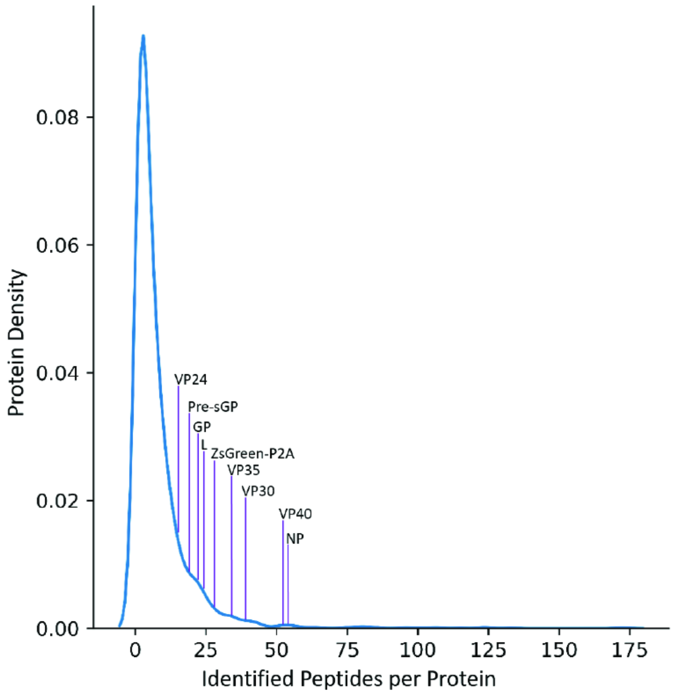


**b**

| Viral Protein Identified | Peptides Identified from Viral Protein |
|---|---|
| NP | 54 |
| VP35 | 34 |
| ZsGreen-P2A | 28 |
| VP40 | 52 |
| Pre-sGP | 19 |
| GP | 22 |
| VP30 | 39 |
| VP24 | 15 |
| L | 24 |

**Supplementary Figure 2**

**Density plot of all proteins identified by mass spectrometry of SuBK12-08 cells infected with LLOV-ZsG-IR$_{ins}$-EBOV$_{UTR+tr}$ (rLLOV$_{comp}$).** (**a**) Kernel density plot indicating the distribution of the number of identified peptides per protein across all 3,939 identified proteins within the rLLOV$_{comp}$-infected SuBK12-08 samples described in Fig 1g. The plot is annotated with the location of all viral proteins within the distribution. (**b**) Table of rLLOV$_{comp}$-encoded proteins and the number of peptides detected by mass spectrometry.

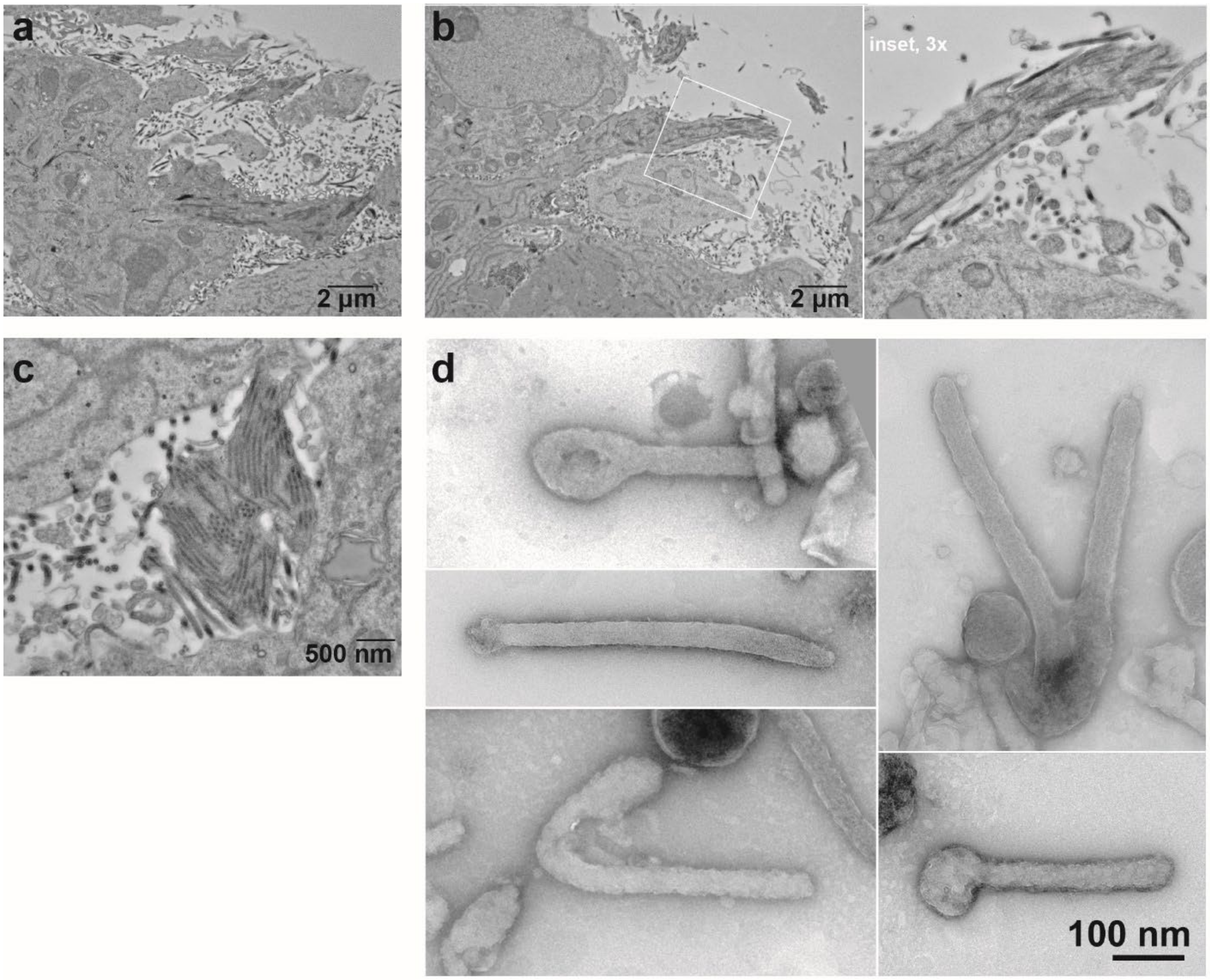


**Supplementary Figure 3**
**EBOV-infected cells and virions.** Huh7 cells were infected with EBOV at an MOI of 5 and fixed at 2 dpi. (**a**) Thin section of EBOV-infected cell releasing viral particles. (**b**) Thin section shows virus budding; inset shown on the right is magnified 3x; protein is black. (**c**) Circled area indicates EBOV inclusions containing cross sectioned and longitudinal sectioned nucleocapsids. (**d**) Negatively stained, isolated EBOV virions show heterogeneous ultrastructure, including 6-shaped, filamentous, shepherd's hook, and branched filamentous structures, as seen previously [59]; protein is white.

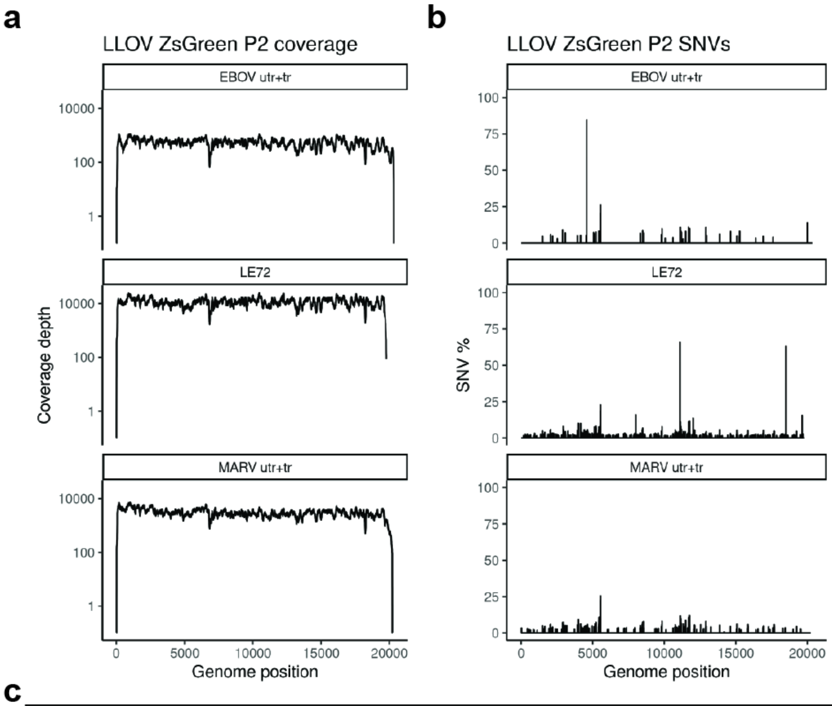


**c**

| Virus | Position | Percent | Reference | SNV | Annotation | Region |
|---|---|---|---|---|---|---|
| EBOV$_{UTR+tr}$ | 4577 | 84.7 | A | ATT | Noncoding A4577ATT | VP35-VP40 IR |
| EBOV$_{UTR+tr}$ | 5533 | 20.5 | A | C | ZsGreen-P2A S240R | P2A |
| EBOV$_{UTR+tr}$ | 5534 | 13.6 | G | C | ZsGreen-P2A S240T | P2A |
| EBOV$_{UTR+tr}$ | 5550 | 26.1 | T | G | ZsGreen-P2A T5550G | P2A |
| EBOV$_{UTR+tr}$ | 11133 | 10.6 | A | C | Noncoding A11133C | VP30-VP24 IR |
| EBOV$_{UTR+tr}$ | 11694 | 10.8 | A | C | Hypothetical F72V | VP30-VP24 IR / rcORF |
| EBOV$_{UTR+tr}$ | 12911 | 10.5 | A | C | Noncoding A12911C | VP24-L IR |
| EBOV$_{UTR+tr}$ | 19999 | 13.7 | GT | G | Noncoding GT19999G | EBOV trailer |
| MARV$_{UTR+tr}$ | 5438 | 10.7 | A | C | ZsGreen-P2A D208A | ZsGreen |
| MARV$_{UTR+tr}$ | 5533 | 22.5 | A | C | ZsGreen-P2A S240R | P2A |
| MARV$_{UTR+tr}$ | 5534 | 13.7 | G | C | ZsGreen-P2A S240T | P2A |
| MARV$_{UTR+tr}$ | 5537 | 11.6 | T | G | ZsGreen-P2A L241R | P2A |
| MARV$_{UTR+tr}$ | 5540 | 11.2 | T | G | ZsGreen-P2A L242R | P2A |
| MARV$_{UTR+tr}$ | 5546 | 14.7 | A | C | ZsGreen-P2A Q244P | P2A |
| MARV$_{UTR+tr}$ | 5550 | 25.5 | T | G | ZsGreen-P2A T5550G | P2A |
| MARV$_{UTR+tr}$ | 11133 | 11.6 | A | C | Noncoding A11133C | VP30-VP24 IR |
| MARV$_{UTR+tr}$ | 11694 | 10.5 | A | C | Hypothetical F72V | VP30-VP24 IR / rcORF |
| MARV$_{UTR+tr}$ | 11770 | 11.9 | A | C | Hypothetical C46W | VP30-VP24 IR / rcORF |
| LE$_{72}$ | 4002 | 10 | T | A | VP35 N297K | VP35 |
| LE$_{72}$ | 4149 | 10 | C | CA | Noncoding C4149CA | VP35-VP40 IR |
| LE$_{72}$ | 5533 | 18.7 | A | C | ZsGreen-P2A S240R | P2A |
| LE$_{72}$ | 5534 | 11.8 | G | C | ZsGreen-P2A S240T | P2A |
| LE$_{72}$ | 5537 | 10.4 | T | G | ZsGreen-P2A L241R | P2A |
| LE$_{72}$ | 5546 | 12.2 | A | C | ZsGreen-P2A Q244P | P2A |
| LE$_{72}$ | 5550 | 22.7 | T | G | ZsGreen-P2A T5550G | P2A |
| LE$_{72}$ | 8012 | 15.9 | T | G | GP1 T8012G | GP1 |
| LE$_{72}$ | 11100 | 65.7 | C | T | Noncoding C11100T | VP30-VP24 IR |
| LE$_{72}$ | 11133 | 11 | A | C | Noncoding A11133C | VP30-VP24 IR |
| LE$_{72}$ | 11694 | 11 | A | C | Hypothetical F72V | VP30-VP24 IR / rcORF |
| LE$_{72}$ | 11770 | 11.5 | A | C | Hypothetical C46W | VP30-VP24 IR / rcORF |
| LE$_{72}$ | 12023 | 13.5 | G | A | VP24 V49I | VP24 |
| LE$_{72}$ | 18494 | 63 | C | T | L A1827V | L |
| LE$_{72}$ | 19638 | 15.6 | C | T | Noncoding C19638T | LLOV L 3' UTR |

**Supplementary Figure 4**

**RNA-Seq data from rLLOV$_{comp}$ clones.** (**a**) Total RNA sequencing of passage 2 rLLOV$_{comp}$ stocks yielded viral genome coverage ≥ 100x with the exception of the leader and trailer regions. Single nucleotide variable (SNV) plot (**b**) and table (**c**), indicating the abundance and location within the genome of SNVs (SNVs ≥ 10% included in the table). Few consensus SNVs (> 50%) were observed, whereas multiple subconsensus SNVs were observed at 10-20%. Mutations within the reporter gene (ZsGreen-P2A) are indicated in green.

a

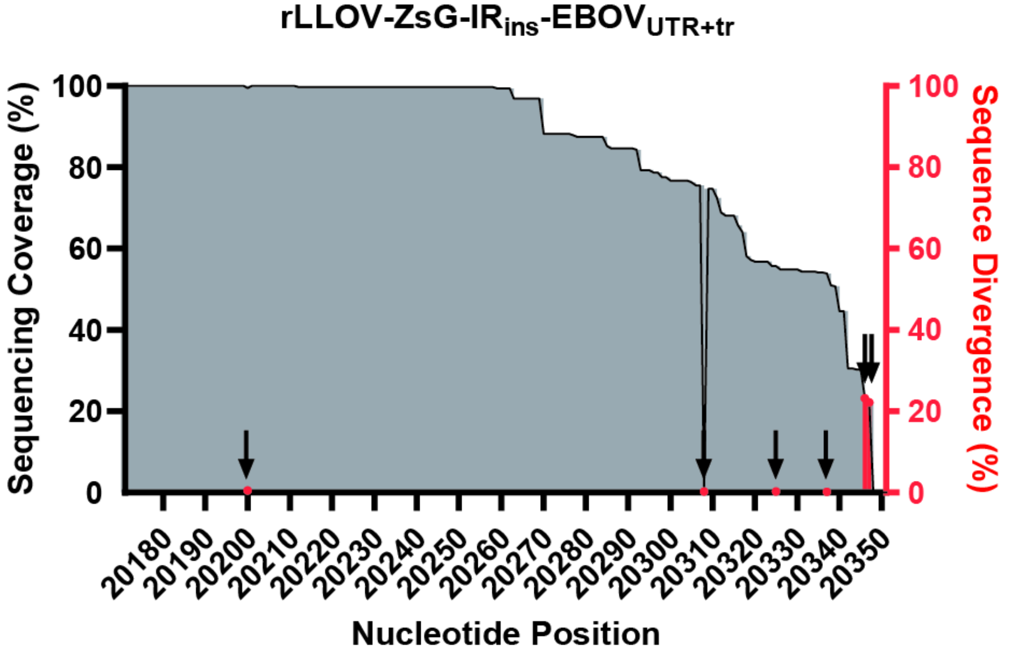


b

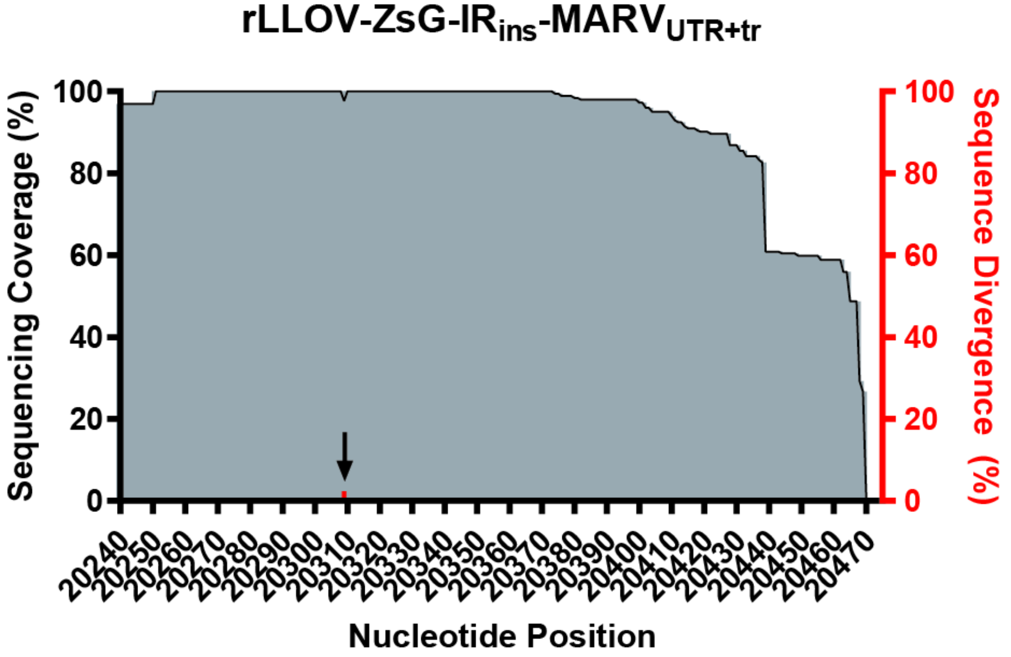


**Supplementary Figure 5**

**Sequence coverage and divergence of rLLOV$_{comp}$ 5' genome ends.** Sequence coverage of the trailer region of both (**a**) rLLOV-EBOV$_{UTR+tr}$ and (**b**) rLLOV-MARV$_{UTR+tr}$ virus stocks. Sequencing coverage percentages were calculated from the number of reads mapping to individual nucleotides within the respective viral genomes. Divergence from the EBOV trailer or MARV trailer sequences are plotted in red and highlighted with a black arrow. Sequencing results captured the terminal 175 nucleotides of the rLLOV-EBOV$_{UTR+tr}$ sequence and the terminal 230 nucleotides of the rLLOV-MARV$_{UTR+tr}$ sequence. Very few differences were observed between the rLLOV$_{comp}$ full-length clone sequences and the viral stocks (passage 2). Very low abundance differences identified in the rLLOV-EBOV$_{UTR+tr}$ stock include a deletion (position 20200, found in 0.58% of reads), an A insertion (between nucleotides 20308 and 20309, 0.33% of reads and the cause of the observed large dip in the graph due to its low abundance), a U to A mutation (position 20325, 0.3% of reads), and a U deletion (position 20337, 0.24% of reads) all in distinct, truncated species with very low read abundance. A significant portion of the reads (22.21% of reads) contained a terminal AC dinucleotide which is not present within the corresponding full-length clone. Similar to the data for the rLLOV-EBOV$_{UTR+tr}$ stock, there are few differences noted between the rLLOV-MARV$_{UTR+tr}$ sequence and corresponding rLLOV$_{comp}$ full-length clone. The only low abundance difference observed was a single A deletion (position 20309, 2.35% of reads). There were no terminal nucleotide additions present in any of the rLLOV-MARV$_{UTR+tr}$ sequence reads.

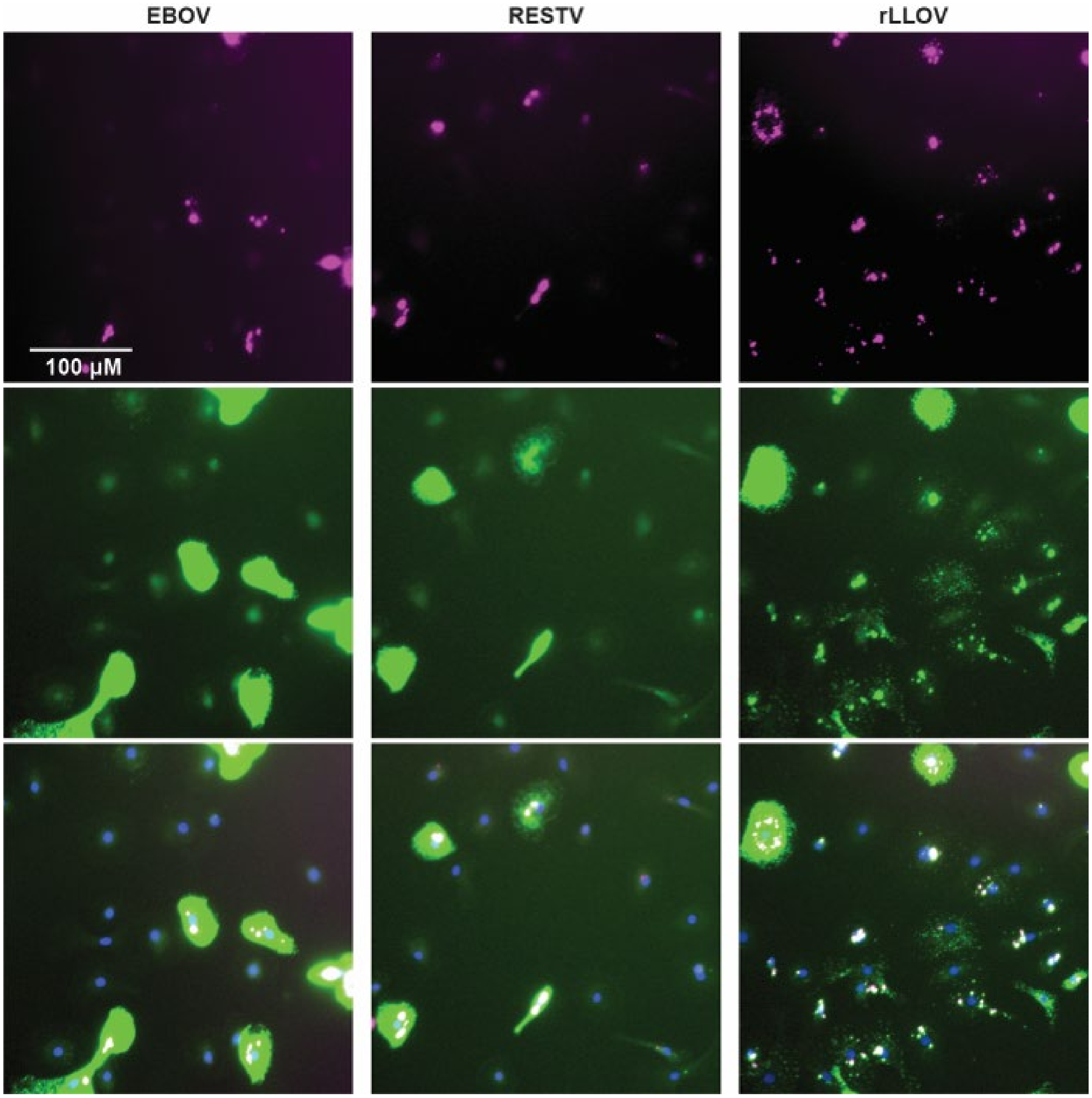


**Supplementary Figure 6**

**Supplementary figure to Fig 4: RNA FISH analysis of human monocyte-derived macrophages (MDMs) infected with EBOV, RESTV, or rLLOV**. MDMs derived from donor 1 were infected at an MOI of 10 with EBOV, RESTV, or rLLOV-IR$_{mut}$-MARV$_{UTR+tr}$. At 1 dpi, cells were fixed and analyzed for the presence of negative (genome, magenta, in top row) and positive (mRNA and antigenome, green, in middle row) sense viral RNA by RNA FISH. Cells were co-stained with DAPI (shown with merged channels, in bottom row). 20x images are shown with increased brightness in green and magenta channels to clearly visualize lower-intensity staining observed in rLLOV-infected cells.
